# Beyond alignment: synergistic integration is required for multimodal cell foundation models

**DOI:** 10.64898/2026.02.23.707420

**Authors:** Till Richter, Eric Zimmermann, James Hall, Fabian J. Theis, Srivatsan Raghavan, Peter S. Winter, Ava P. Amini, Lorin Crawford

## Abstract

The vision of a “virtual cell”—a computational model that simulates biological function across modalities and scales—has become a defining goal in computational biology. While powerful unimodal foundation models exist, the lack of large-scale paired data prohibits the joint training of multimodal approaches. This scarcity favors compositional foundation models (CFMs): architectures that fuse frozen unimodal experts via a learned interface. However, it remains unclear when this multimodal fusion adds task-relevant information beyond the strongest unimodal representation and when it merely aggregates redundant signal. Here, we introduce the Synergistic Information Score (SIS), a metric grounded in partial information decomposition (PID), that quantifies the information gain achievable only through cross-modal interactions. Extending theoretical results from self-supervised learning, we show that standard alignment-based fusion objectives on frozen encoders inherently collapse to detecting linear redundancies, limiting their ability to capture nonlinear synergistic states. This distinction is directly relevant for tasks aiming to link tissue morphology and gene expression. Benchmarking ten fusion methods on spatial transcriptomics datasets, we use SIS to demonstrate that tasks dominated by linear redundancies are sufficiently served by unimodal baselines, whereas complex niche definitions benefit from synergy-aware integration objectives that enable cross-modal interactions beyond linear alignment. Finally, we perform a scaling analysis which highlights that fine-tuning a dominant unimodal expert is the most sample-efficient path for standard tasks, suggesting that the benefits of multimodal frameworks only emerge when tasks depend on information distributed across modalities. Together, these results establish that building towards a virtual cell will require a fundamental shift from alignment objectives that emphasize shared structure to synergy-maximizing integration that preserves and exploits complementary cross-modal signal.

## Introduction

Advances in self-supervised learning and the increasing scale of biological datasets have produced powerful encoders for individual modalities, including transcriptomics [1–4], histopathology [5–7], and spatial transcriptomics [8–10]. Recent multimodal efforts extend these representations to chromatin accessibility [11, 12], proteomics [13], and text-based biological metadata [14–17]. However, multimodal data remains fundamentally asymmetric: millions of unimodal samples exist in isolation, whereas paired measurements capturing multiple modalities in the same cell or tissue context are significantly less available. This asymmetry favors compositional foundation models (CFMs), which reuse robust unimodal encoders as frozen experts and combine their outputs through learned interfaces [18]. These interfaces enable scalable multimodal modeling without requiring end-to-end retraining and have become central to efforts aimed at constructing integrated representations of cellular state, an essential step toward “virtual cell” models that integrate multiple modalities into unified representations of cellular identity and function [19–21] (see discussion on related work in Supplementary Note 1). Still, a fundamental question remains: when does multimodal fusion truly add information beyond the strongest unimodal expert, and when does it merely repackage redundant signal?

Standard alignment-based fusion interfaces, which learn mappings between frozen unimodal representations into a shared embedding space (formally defined in Methods), are optimized to maximize agreement between modalities. This is typically done by bringing representations of paired samples closer while separating unrelated samples through contrastive or regression-based objectives [22, 23]. These objectives are effective for retrieval and matching tasks, where the goal is to collapse modalities onto a shared structure (e.g., CLIP [22]). However, we argue that many scientific objectives require the opposite regime where discovery arises from synthesizing non-overlapping sets of information. For example, defining spatially varying cell states, niches, or microenvironmental interactions may benefit from fusing tissue morphology and gene expression. In these regimes, we hypothesize that alignment-based fusion may reach a spectral ceiling defined here as a limit imposed by the eigenspectrum of pretrained representations, such that alignment recovers dominant linear correlations present in pretrained encoders but may fail to access synergistic information that emerges only through joint nonlinear processing. We investigate this limitation both theoretically and empirically.

In this work, we formalize and assess the limits of alignment-based CFMs under a central practical constraint of multimodal biology: unimodal datasets are abundant, whereas high-quality paired measurements remain comparatively scarce [19, 21]. At the same time, integrating complementary modalities is increasingly recognized as essential for capturing cellular state, motivating accelerated investment in multimodal data generation and modeling [9, 18, 24]. As a result, many fusion strategies must operate in low-pairing regimes, where interfaces are trained with limited paired supervision. We demonstrate that simple alignment can *appear* effective while still primarily propagating redundant structure and suppressing modality-specific signal—a failure mode that is difficult to detect from downstream performance alone without an explicit diagnostic.

Our study can be broken down into three main components:

- First, we introduce the Synergistic Information Score (SIS), a diagnostic grounded in partial information decomposition (PID) [25, 26], which quantifies whether a downstream task truly benefits from multimodal integration beyond the strongest unimodal expert. SIS measures the relative performance gain a fused representation provides over the best unimodal representation under a fixed probe class (here, linear probes), and thus it captures when fusion makes additional task-relevant information accessible rather than merely aggregating redundant signal. In other wo158.rds, SIS serves as a diagnostic to determine whether a task is *unimodal-sufficient* or *cross-modal-dependent* (Fig. 1).
- Second, we use SIS to show that many frozen-encoder alignment objectives saturate because they learn linear redundancy (i.e., hit a spectral ceiling), which causes them to fail on cross-modal-dependent tasks. To explain this behavior, we take spectral theory from the self-supervised learning literature and expand it to the multimodal frozen encoder setting [23, 27]. Our extension shows why diverse alignment objectives converge to equivalent spectral solutions that optimally recover redundancy but are limited in their ability to access nonlinear synergistic information.
- Third, we evaluate these predictions using a comprehensive experimental suite across multiple spatial transcriptomics datasets. Specifically, our evaluation uses a pulmonary fibrosis lung dataset [28, 29], a developmental human thymus dataset [30, 31], and a breast cancer dataset [32, 33] (see Supplementary Note 2 for dataset details). These datasets span regimes ranging from tight correspondence between modalities to substantial resolution mismatch. SIS separates regimes where unimodal finetuning suffices to extract all task-relevant information accessible to linear probes from regimes where multimodal integration provides additional predictive signal, exposing the value of paired data when modeling biology.

**Figure 1.**
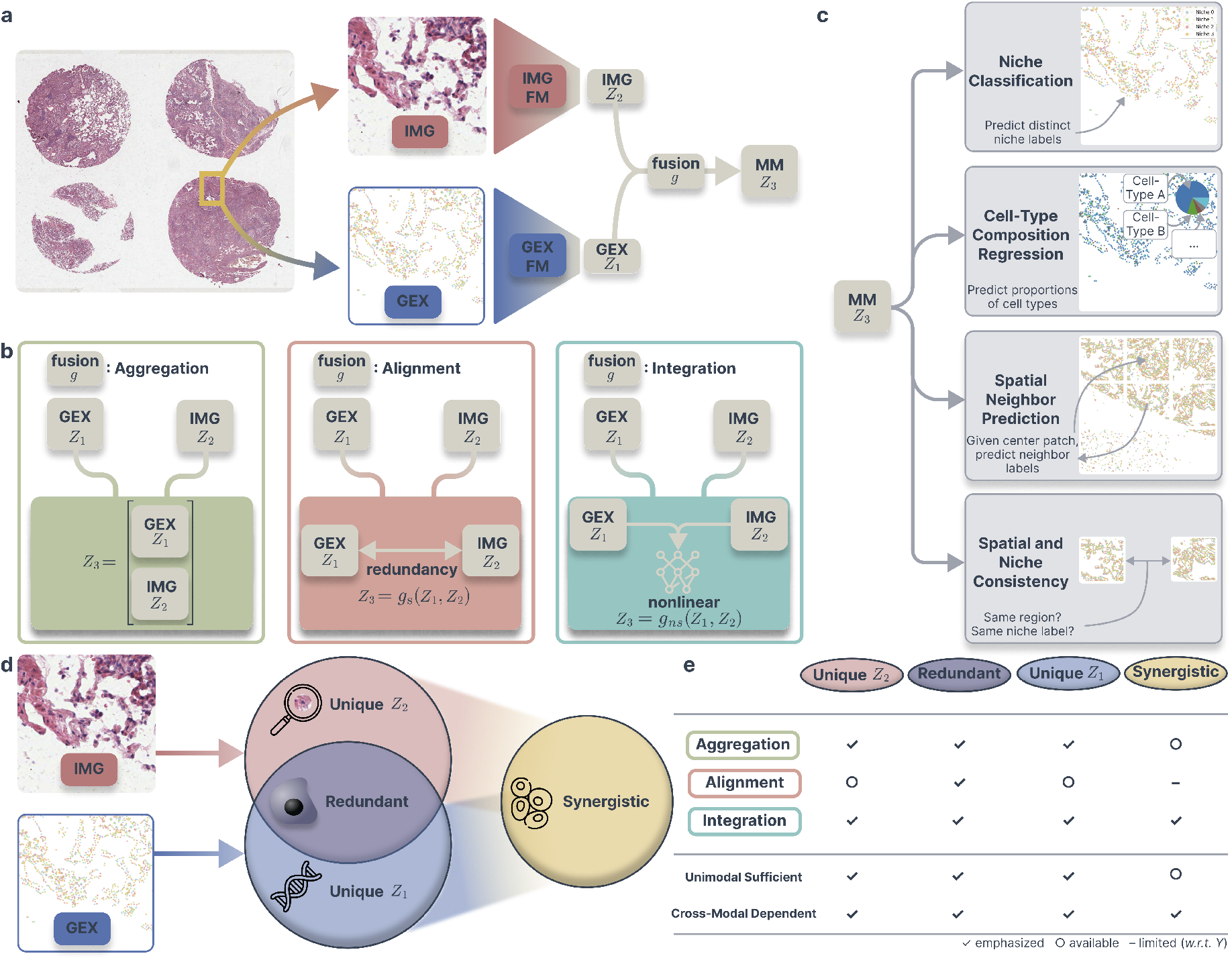
Fusion in compositional foundation models. **a**, Schematic of the system architecture. We freeze unimodal foundation models (for histopathology images and gene expression, IMG and GEX respectively) and train only the fusion interfaces *g*_1_ and *g*_2_ (summarized as *g*; details in Methods); downstream evaluation uses the multimodal (MM) concatenation of fused representations (*z*_3_) and a linear probe. This isolates the fusion mechanism as the sole variable contributing to the evaluation. **b**, Schematic of fusion methods: aggregation (left), spectral (middle), non-spectral (right). **c**, Downstream evaluation tasks probing different information regimes. We evaluate fused embeddings on niche classification, cell-type composition regression, spatial neighborhood prediction, and spatial and niche consistency. **d**, Conceptual decomposition of multimodal biological information into unique, redundant, and synergistic components with respect to a downstream target *Y* (represented abstractly by the prediction tasks in panel c). Unique components are modality-specific, redundant components are shared across modalities, and synergistic components arise only through joint reasoning over modalities. **e**, Information-theoretic view of multimodal fusion. Different fusion strategies emphasize different information components. Symbols indicate whether unique, redundant, or synergistic information (with respect to the target *Y*) is emphasized (✓), available (◦), or limited (−). This panel highlights which information components are emphasized, available, or limited for each fusion strategy and downstream task.

Beyond standard downstream tasks such as niche classification and cell-type composition regression, we evaluate spatial tasks that probe spatial understanding, including niche consistency, spatial consistency, and neighbor prediction. These tasks span both unimodal-sufficient regimes, where transcriptomic information alone supports accurate prediction, and cross-modal-dependent regimes, where integrating morphology improves performance. We further perform a fine-tuning scaling analysis to characterize how the value of multimodal integration changes with dataset size. Across our benchmarks, synergy-aware objectives (e.g., CoMM [34]) can break the spectral ceiling in cross-modal-dependent regimes, whereas alignment-based methods typically saturate near unimodal performance.

Together, our analyses establish SIS as a diagnostic of the information regime that governs if and how a given task benefits from multimodal learning. We show that when tasks are unimodal-sufficient, multimodal fusion provides limited additive benefit beyond strengthening the dominant modality; however, when tasks are cross-modal-dependent, integration can expose complementary signal that is inaccessible to unimodal representations alone. This distinction serves as a principled guide to whether improvements in predictive performance reflect unimodal refinement or genuine benefit from cross-modal pairing. More broadly, it also clarifies when multimodal foundation models can support truly synergistic inference—moving beyond correspondence to biological synthesis across scales, as envisioned by the synthesis-oriented objectives of virtual cell efforts.

## Results

### Multimodal integration in compositional cell foundation models

The scarcity of paired multimodal data limits end-to-end training of envisioned virtual cells. This scarcity favors compositional foundation models (CFMs): architectures that combine robust, pretrained unimodal experts via a lightweight learned fusion interface. In this framework, the foundation model encoders are frozen, and the multimodal synthesis is entirely based on the fusion mechanism. To systematically evaluate these mechanisms, we established a controlled benchmarking framework (Fig. 1).

Our experimental design (Fig. 1a) rigorously isolates the contribution of the fusion strategy. Specifically, we extract spatially matched patches from whole-slide histopathology and spatial gene expression data (Supplementary Note 2), ensuring that both modalities view the same physical tissue region. We employ state-of-the-art frozen foundation model encoders, using UNI-2 [5] for histopathology and Nicheformer [8] for spatial transcriptomics, to transform matched patches from the same tissue slide into latent embeddings (Methods and Supplementary Note 3). Because the foundation model encoders remain fixed, any difference in downstream performance is attributable solely to the fusion interface. We benchmark ten distinct fusion interfaces, ranging from simple aggregation (e.g., Concatenation) to spectral (e.g., Canonical-Correlation Analysis (CCA) and VICReg [35]) and non-spectral objectives (e.g., CoMM [34]) (Fig. 1b and Supplementary Notes 4-5).

Standard benchmarks often obscure the distinction between simple redundancy and genuine multimodal integration. To resolve this, we design a set of evaluations that probe distinct regimes of structural dependency between modalities (Fig. 1c). Specifically, we consider three primary downstream tasks that span increasing spatial and semantic complexity:

- **Local phenotyping evaluations**. *Cell-type composition regression* and *niche classification* primarily probe information available within a local patch, and therefore likely reside in a unimodal-sufficient regime.
- **Spatial structuring evaluation**. *Spatial neighborhood prediction* requires inferring tissue organization beyond the local field of view, potentially increasing reliance on cross-modal information.

In addition to these primary tasks, we also evaluate *spatial* and *niche consistency* as auxiliary probes, assessing whether the models capture spatial information encoded by spatial location and niche label (see Methods). By evaluating methods across this spectrum, we assess *when* alignment works, distinguishing unimodal-sufficient regimes, where unimodal representations already capture task-relevant information, from cross-modal-dependent regimes, where multimodal integration provides additional predictive signal.

### The Synergistic Information Score distinguishes integration from alignment

A core question in evaluating CFMs is whether multimodal predictions benefit from genuinely complementary information or merely from aggregating redundant features across views. Standard downstream metrics such as F1 scores or R-squared (*R*^2^) quantify predictive accuracy, but are agnostic to the information structure underlying that accuracy, specifically in distinguishing whether improvements stem from shared, unique, or synergistic information between modalities. This distinction is central in biological settings, where cells are measured through multiple complementary views (e.g., morphology and gene expression), and where differences between measurements may either be noise to suppress or signal to exploit (Fig. 1d).

To make this distinction explicit, we introduce the Synergistic Information Score (SIS), a metric grounded in partial information decomposition (PID) and that isolates information about a target *Y* that is accessible *only through cross-modal interaction* (Methods). SIS operationalizes the conceptual dichotomy illustrated in Fig. 1: *alignment* treats two measurements of the same entity as partially redundant views whose differences should be ignored, whereas *integration* treats those differences as informative and potentially interacting.

Formally, let *Z*_1_ and *Z*_2_ denote frozen unimodal representations, and let *Z*_3_ = [*g*_1_(*Z*_1_); *g*_2_(*Z*_2_)] be their fused representation produced by a fusion interface 𝒢 = (*g*_1_, *g*_2_). Given a downstream task variable *Y* (e.g., niche labels), we use the minimum mutual information assumption [25, 26] to define SIS as the relative information gain beyond the strongest unimodal baseline:

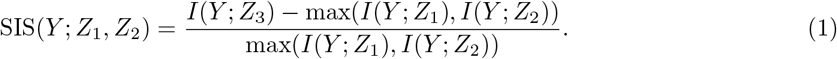

In practice, the intractable mutual information *I*(*Y* ; *Z*) is estimated via linear probes trained on the frozen representations (i.e., *Z*_1_ and *Z*_2_ in the unimodal case, and *Z*_3_ in the multimodal case). For classification tasks we use F1 Macro scores, and for regression tasks we use *R*^2^. Restricting our evaluation to linear probes is crucial: SIS measures whether a fusion interface has rendered synergistic information *linearly accessible*, rather than deferring integration to complex downstream nonlinear models. Positive SIS indicates accessible synergy; near-zero or negative SIS indicates redundancy dominance or fusion failure. Box 1 summarizes a practical workflow and links SIS to tasks that are unimodal sufficient versus those that are cross-modal dependent (Fig. 1e).

Evaluating ten fusion strategies across three spatial transcriptomic datasets (a pulmonary fibrosis lung atlas [28, 29], a developmental human thymus dataset [30, 31], and a breast cancer dataset [32, 33]; Supplementary Figs. 1-3) reveals that multimodal fusion is inherently task-dependent (Table 1). Consistent with the taxonomy in Fig. 1e, spectral alignment approaches emphasize shared information, aggregation methods preserves modality-specific unique information, and non-spectral integration methods enable synergistic interactions. We next asked when multimodal fusion is helpful in the frozen-encoder setting, and when a single modality suffices. To this end, SIS serves as a diagnostic of whether a given biological task benefits from multimodal learning, or whether performance is already saturated by a unimodal expert.

**Table 1.**
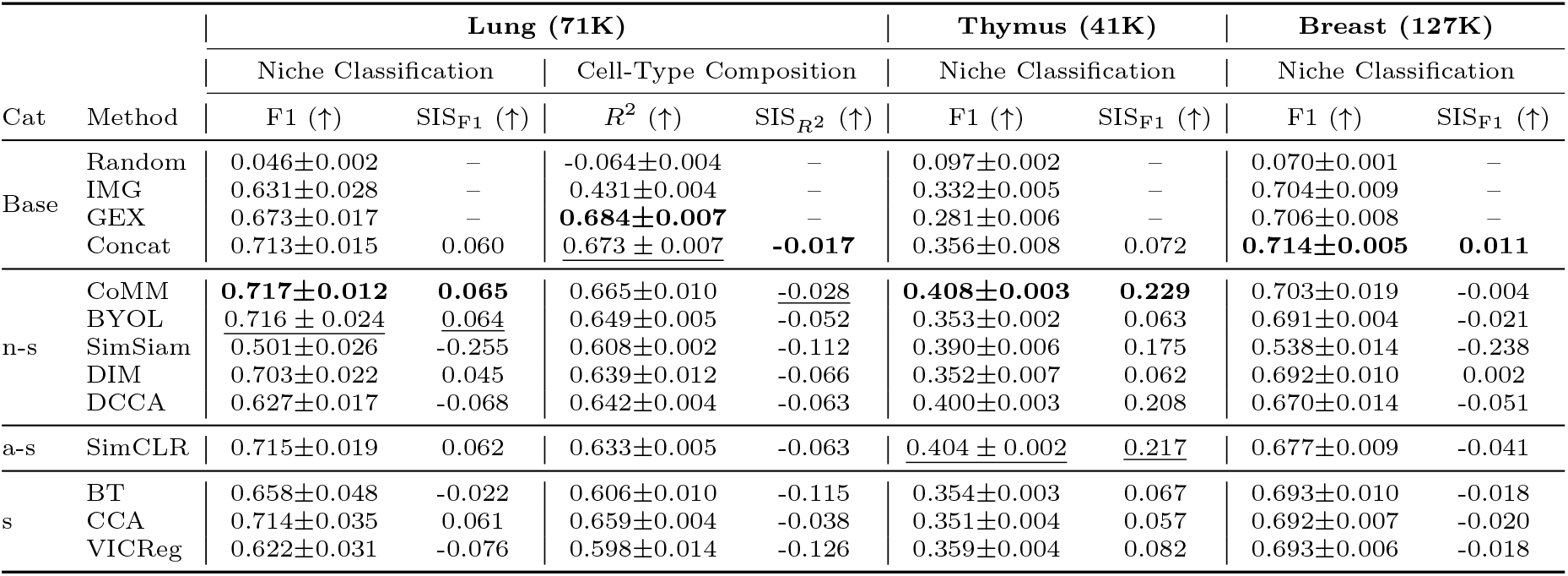
Task-dependent multimodal synergy across spatial transcriptomic datasets. Columns report the niche classification F1 scores and SIS based on F1 (denoted as SIS_F1_) for the lung, thymus, and breast datasets. For the lung dataset, we also report the *R*^2^ results for the cell-type composition regression task and the corresponding SIS based on *R*^2^ (denoted as 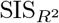). Note that SIS_F1_ and 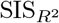 are task-specific and reported per downstream metric. Values are the mean ± standard deviation across seeds; dashes indicate unavailable metrics. The best performing method for each task is listed in **bold**, and the second best method is underlined. Numbers in parentheses (71K, 41K, and 127K) indicate the total number of spatially matched patches used for evaluation (see Supplementary Table 1 for details). Method categories include: “base” (unimodal or trivial baselines), “s” (spectral alignment methods reducing to linear alignment), “a-s” (approximately spectral methods converging to the spectral solution in limiting regimes), and “n-s” (non-spectral integration methods introducing learned nonlinear interactions). Definitions and justification are given in Methods. IMG = imaging, GEX = gene expression, and Concat = Concatenation.

#### Box 1

**Using the Synergistic Information Score (SIS) in practice**.

**SIS as a diagnostic**. SIS quantifies the performance gain of a fused representation relative to the *best* unimodal representation under a fixed probe class (here, linear probes). SIS distinguishes whether a task is *unimodal-sufficient* (reflected by SIS ≲ 0), where one modality already captures the relevant signal, or *cross-modal-dependent* (SIS > 0), where integrating modalities makes additional task-relevant information accessible.

**Recommended workflow**.

1. **Evaluate on paired data**. On a representative paired subset, train unimodal probe(s) and a fused probe, and compute SIS.
2. **Align modeling and data strategy with the information regime**. For unimodalsufficient tasks (SIS ≲ 0), improvements typically follow from strengthening the dominant modality. For example, this could be through higher-quality unimodal data, improved labels, or targeted fine-tuning. For cross-modal-dependent tasks (SIS > 0), multimodal integration can expose complementary signal not available in isolation, motivating investment in paired measurements and integration-aware interfaces. This regime is desired in synthesis-oriented objectives, such as efforts envisioning “virtual cells” which aim to combine modalities into a unified representation of cellular state.
3. **Probe robustness**. Recompute SIS across task variants that modulate ambiguity or context (e.g., increasing spatial distance or neighborhood size). Increasing SIS indicates growing reliance on cross-modal interaction.

**Interpretation**. SIS is probe-dependent and, in this work, measures *linearly accessible* synergy. It should be interpreted as a diagnostic of the task’s information regime rather than a prescriptive criterion: low SIS indicates that fusion does not improve accessibility of task-relevant signal under the current setup, whereas positive SIS identifies regimes where integration provides measurable benefit.

### Alignment can destroy unique information

Task dependence is most evident in the breast dataset, which consists of large tissue patches with dense cellular content. Here, for the niche classification task, the naive Concatenation baseline achieves the highest SIS (SIS = 0.011; Table 1), outperforming more complex integration methods such as CoMM [34] (SIS = -0.004) and SimCLR [36] (SIS = -0.041). This exposes a trade-off implicit in many multimodal objectives: alignment-based methods (e.g., CCA, contrastive objectives) explicitly encourage *g*_1_(*Z*_1_) ≈ *g*_2_(*Z*_2_), thereby suppressing modality-specific signals that do not correlate across views. When a task relies on independent, complementary cues from each modality, this suppression is harmful. Concatenation, by preserving unique information, is better matched to this regime, and its positive SIS on the breast dataset confirms that alignment can be counterproductive when differences between views are informative. Similar behavior is observed in lung niche classification, where Concatenation achieves comparable SIS to alignment and integration methods (Table 1).

#### Synergy is required to integrate modalities

Aggregation fails when a task depends on integrating complementary information across modalities. In the thymus dataset, gene expression is measured using Visium spatial transcriptomics [30, 37], which captures transcripts over coarse spots spanning multiple cells and therefore does not align precisely with the fine-grained structure visible in histology images. This known resolution mismatch creates an integration regime where complementary spatial information is distributed across modalities. In this setting, Concatenation yields low synergy (SIS = 0.072) among benchmarked fusion methods, reflecting its inability to reconcile spatial mismatch (Table 1). By contrast, the synergy-aware method CoMM achieves substantially higher synergy (SIS = 0.229), exceeding alignment baselines such as CCA (SIS = 0.059) by a wide margin, demonstrating that the task is cross-modal dependent and requires integration rather than simple aggregation. This demonstrates that when correspondence across views is imperfect, effective fusion requires learning nonlinear cross-modal interactions rather than enforcing alignment. A similar but weaker pattern appears in the lung niche classification task (Table 1), where spatial correspondence is tighter due to higher-resolution spatial transcriptomics measurements. In this regime, aggregation and alignment perform closer to integration methods, indicating reduced, though not eliminated, dependence on cross-modal interaction.

#### Unimodal dominance renders integration unnecessary

The lung cell-type regression task provides a control regime where unimodal gene expression signal is known to be highly informative. Gene expression is a primary molecular determinant of cell identity, as established by large-scale single-cell atlases and transcriptomic profiling studies [38, 39]. As a result, for this downstream task, we reasoned that histology would provide information that is redundant to what is strongly conveyed through gene expression. Consistent with this, SIS is negative across all fusion methods (e.g., CoMM scored SIS = -0.028), indicating that multimodal fusion does not provide additional task-relevant information beyond the strongest unimodal representation, which in this task is gene expression (see Table 1 where GEX *R*^2^ = 0.684 is the highest among the unimodal baselines). This demonstrates the diagnostic value of SIS: it identifies regimes where integration is unnecessary because the relevant signal is already accessible through a single modality.

#### From alignment to integration

Together, these results establish a PID-based view of multimodal fusion (Fig. 1d,e). Fusion interfaces applied to frozen unimodal encoders do not uniformly improve performance, but instead reflect the task’s underlying information regime. Spectral alignment methods preferentially extract shared structure but may suppress modality-specific signal; aggregation methods preserve modality-specific information but does not model interactions; and non-spectral integration methods enable access to cross-modal synergy. Consequently, the design of multimodal cell foundation models should be guided not by a single universal fusion strategy, but by the information structure induced by the biological question and downstream task of interest.

### Synergistic integration recovers spatial tissue organization

We next asked how the cross-modal-dependent regime evolves as tasks require increasing spatial context. Spatial context is a defining property of biological tissues: cellular function depends not only on local composition but also on structured interactions across space. Multimodal cell foundation models must therefore support representations that capture tissue-wide organization rather than memorizing local correlations. This capability is reflected in whether models extrapolate to regions physically beyond immediate observation. We hypothesized that, as prediction distance increases and local ambiguity grows, performance should increasingly depend on integrating complementary information across modalities. We probe this capability by measuring SIS in a spatial neighborhood prediction task (Fig. 2a-d), where frozen unimodal encoders and their multimodal fusion are evaluated on predicting niche identity and cell-type composition of neighboring patches at increasing spatial distances (i.e., going from the center to a neighbor patch).

**Figure 2.**
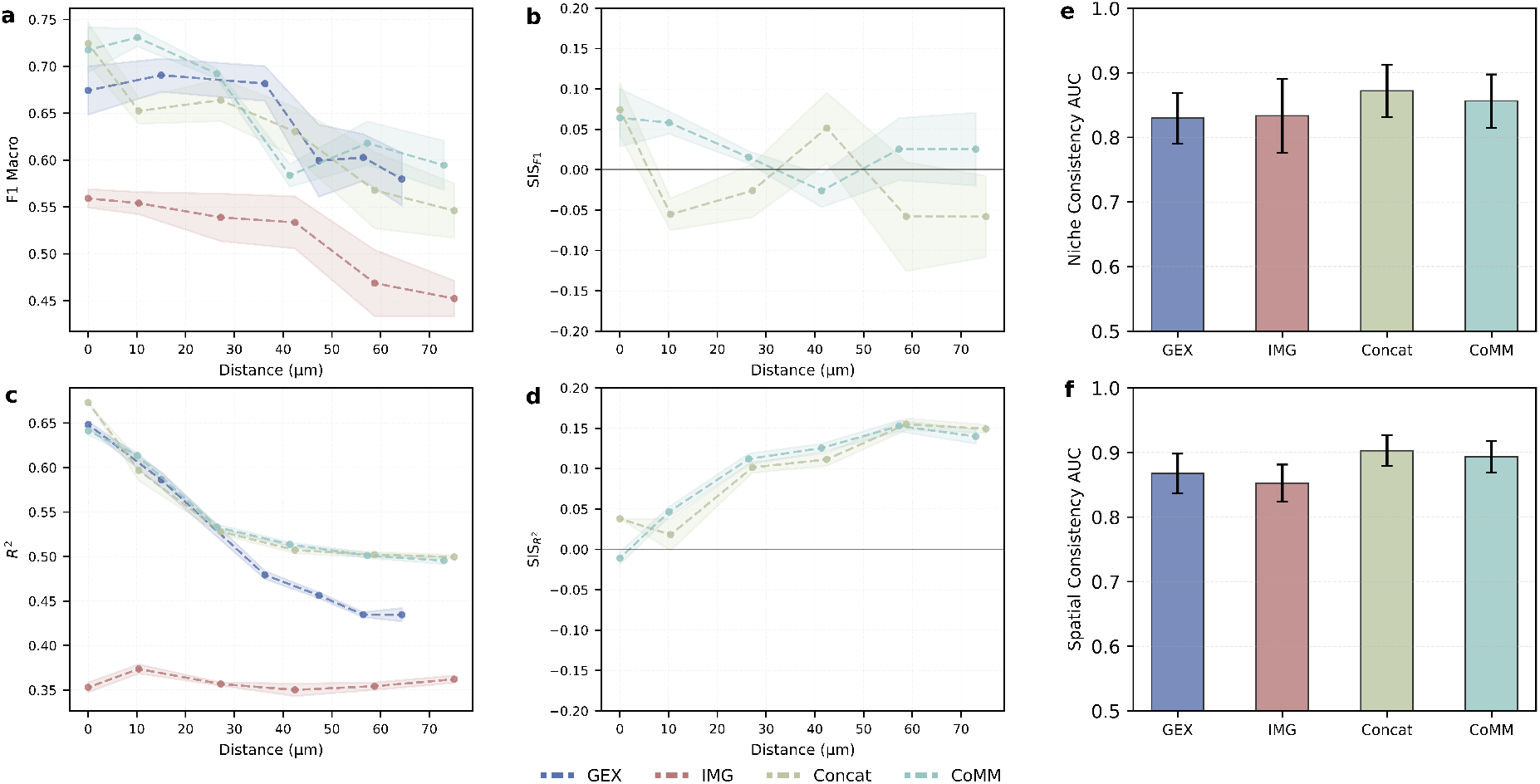
Benchmarking frozen encoders and their fusion on spatial neighborhood tasks in the lung dataset. **a-d**, Spatial neighbor prediction on the lung dataset where performance and corresponding SIS are plotted as a function of the distance between a *center* patch and a *neighbor* patch. Neighbor niche classification is shown in (a) with F1 Macro and in (b) with SIS_F1_. Neighbor cell-type composition regression is shown in (c) with *R*^2^ and in (d) with 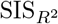. SIS quantifies the performance gain of a fused representation relative to the strongest unimodal representation; the horizontal line marks SIS = 0. In (a) and (c), we show unimodal gene expression (GEX; blue) and imaging (IMG; red) alongside multimodal fusion methods Concat (green) and CoMM (turquoise). In (b) and (d), we report SIS only for the multimodal fusion methods (Concat, CoMM), since SIS is defined relative to the best unimodal baseline. **e-f**, Niche consistency and spatial consistency on the lung dataset, quantified as area under the curve (AUC) for a consistency classifier computed using the similarities between embeddings (Methods). Shaded regions and error bars denote mean ± one standard deviation across five cross-validation folds. Distances with insufficient valid neighbor pairs for a given modality are omitted.

This setting explicitly stress-tests fusion objectives (Fig. 2a-d). At short center-to-neighbor distances, unimodal representations already perform strongly and SIS remains near zero, indicating unimodal-sufficient regimes. As this spatial distance increases, unimodal performance degrades more rapidly than multimodal fusion and SIS correspondingly rises (Fig. 2d), indicating a transition toward a cross-modal-dependent regime where integration becomes increasingly beneficial.

#### Multimodal fusion preserves tissue consistency

We first verify that fusion does not distort the intrinsic biological structure encoded by unimodal experts. To do this, we quantify *niche* and *spatial consistency* via pairwise probes that predict (from two patch embeddings) whether patches share the same niche label or spatial region, summarized by area under the curve (AUC; Methods). Both unimodal encoders exhibit strong spatial and niche consistency (AUC > 0.8; Fig. 2e,f, Supplementary Fig. 4, and Supplementary Note 6), reflecting the strong spatial priors learned during pretraining. Importantly, multimodal fusion preserves this structure across all datasets (Supplementary Fig. 4). Multimodal fusion models achieve consistency comparable to, or marginally exceeding, unimodal baselines, indicating that fusion establishes a shared latent space without collapsing or corrupting the spatial manifold. We additionally verify that fusion interfaces preserve spatial and semantic neighborhood structure via pairwise consistency diagnostics (Supplementary Tables 2-3 and Supplementary Note 7). This confirms that subsequent gains in spatial prediction arise from information synthesis rather than representational degradation.

#### Synergy emerges as spatial context expands

Predicting properties of neighboring patches reveals a clear transition from redundancy-dominated to integration-dominated regimes (Fig. 2a-d). At short center-to-neighbor distances (≤ 10 *µ*m), the neighbor patch remains within the local microenvironment of the center patch, and unimodal signals are relatively predictive (Fig. 2a,c). In this regime, unimodal encoders perform strongly, and multimodal fusion offers only limited improvements over aggregation (Fig. 2a,c), consistent with alignment being sufficient. As spatial distance increases (20–60 *µ*m), local correspondence weakens, and unimodal performance degrades more rapidly than multimodal performance. Notably, in neighbor cell-type composition regression, a continuous task that depends on tissue-scale context rather than discrete local cues, degradation is driven primarily by the gene expression unimodal baseline, while imaging remains comparatively stable (Fig. 2c; some distances are unavailable for specific modalities due to dataset filtering and variable neighbor availability). Accordingly, SIS increases sharply with distance for this task (from ≈ 0 to > 0.15; Fig. 2d), while SIS for neighbor niche classification remains comparatively flat (Fig. 2c). This pattern indicates that as spatial distance grows, predictive success increasingly depends on integrating complementary signals across modalities rather than exploiting shared redundancy.

#### Convergence toward integration under spatial ambiguity

Comparing datasets further highlights how known differences in spatial resolution modulate the utility of fusion (Supplementary Note 6). The lung and breast datasets utilize Xenium *in situ* technology, where gene expression and tissue morphology are measured at comparable effective resolution and exhibit strong local correspondence, resulting in largely redundant representations at short range. Consistent with this, SIS is low or negative at short range, and simple Concatenation performs competitively (Fig. 2b,d and Supplementary Fig. 4a-d In contrast, the thymus dataset uses Visium spatial transcriptomics, which captures gene expression over coarse spots spanning multiple cells and therefore introduces inherent spatial ambiguity relative to high-resolution histology. In this regime, SIS is already positive at short distances for neighbor niche classification (Supplementary Fig. 4f), and the synergy-aware CoMM model consistently outperforms aggregation. Despite these differing starting points, SIS converges to a positive value across all datasets at long range (SIS ≈ 0.03–0.07), indicating a shared setting in which spatial context dominates local redundancy and learned integration becomes essential regardless of initial measurement resolution.

#### When spatial integration becomes necessary

Together, these results show that, for frozen-encoder fusion, the benefit of multimodal integration in neighborhood prediction grows with spatial range and is amplified under resolution mismatch. In regimes with strong local correspondence, fusion provides limited gains over unimodal baselines and simple aggregation; whereas, in regimes with increased spatial ambiguity, integration-aware interfaces yield higher SIS and better preserve performance at larger distances (Fig. 2c,d and Supplementary Fig. 4). This suggests that modeling the emergent structure of tissue biology requires moving beyond objectives that merely align views toward interfaces that actively synthesize complementary spatial information.

### A spectral ceiling constrains frozen alignment

The observation that many fusion methods converge to similar performance plateaus (Table 1) points to a shared theoretical constraint inherent to the frozen encoder setting. Building on self-supervised learning theory showing that diverse unimodal objectives reduce to spectral solutions [23, 27], we extend this analysis to multimodal fusion. Here, the idea of a *spectral ceiling* refers to the constraint that, under frozen encoders and linear fusion mappings with whitening/variance constraints, optimization reduces to an eigen/singular-value problem on the cross-covariance, limiting solutions to linear correlations between modalities. This extension reveals a fundamental distinction: alignment objectives optimize linear redundancy under a spectral ceiling, whereas integration objectives can, in principle, capture nonlinear synergy beyond this limit.

#### Frozen encoders reduce alignment to linear redundancy

In the frozen encoder regime, we show that a broad class of fusion objectives reduces, under linear alignment mappings and standard variance or whitening constraints, to maximizing the trace Tr(*C*), where 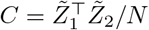 denotes the cross-covariance between aligned representations 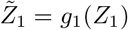 and 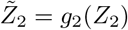 for *N* observations. By the Eckart–Young theorem [40], the optimal solution is given by the singular value decomposition of *C*. In the Methods, we formalize this reduction by showing that, under frozen encoders and linear fusion, a wide class of objectives is equivalent to maximizing Tr(*C*) up to normalization constraints (i.e., whitening during training and *L*_2_-normalization at evaluation). Crucially, the variance or whitening constraints imposed by these objectives are necessary to exclude trivial collapsed solutions (e.g., 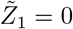), but they also enforce a linear spectral structure. As a result, such methods act as alignment interfaces: they optimally recover linear correlations shared between modalities, but are theoretically restricted to information expressible within the linear subspace of 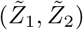. By construction, they cannot exploit the task-relevant structure that appears only through nonlinear cross-modal interactions. In contrast, integration approaches (e.g., CoMM and BYOL) introduce asymmetry or synergy-aware terms that prevent reduction to this eigenvalue problem, allowing them to escape the spectral ceiling (Fig. 3a).

**Figure 3.**
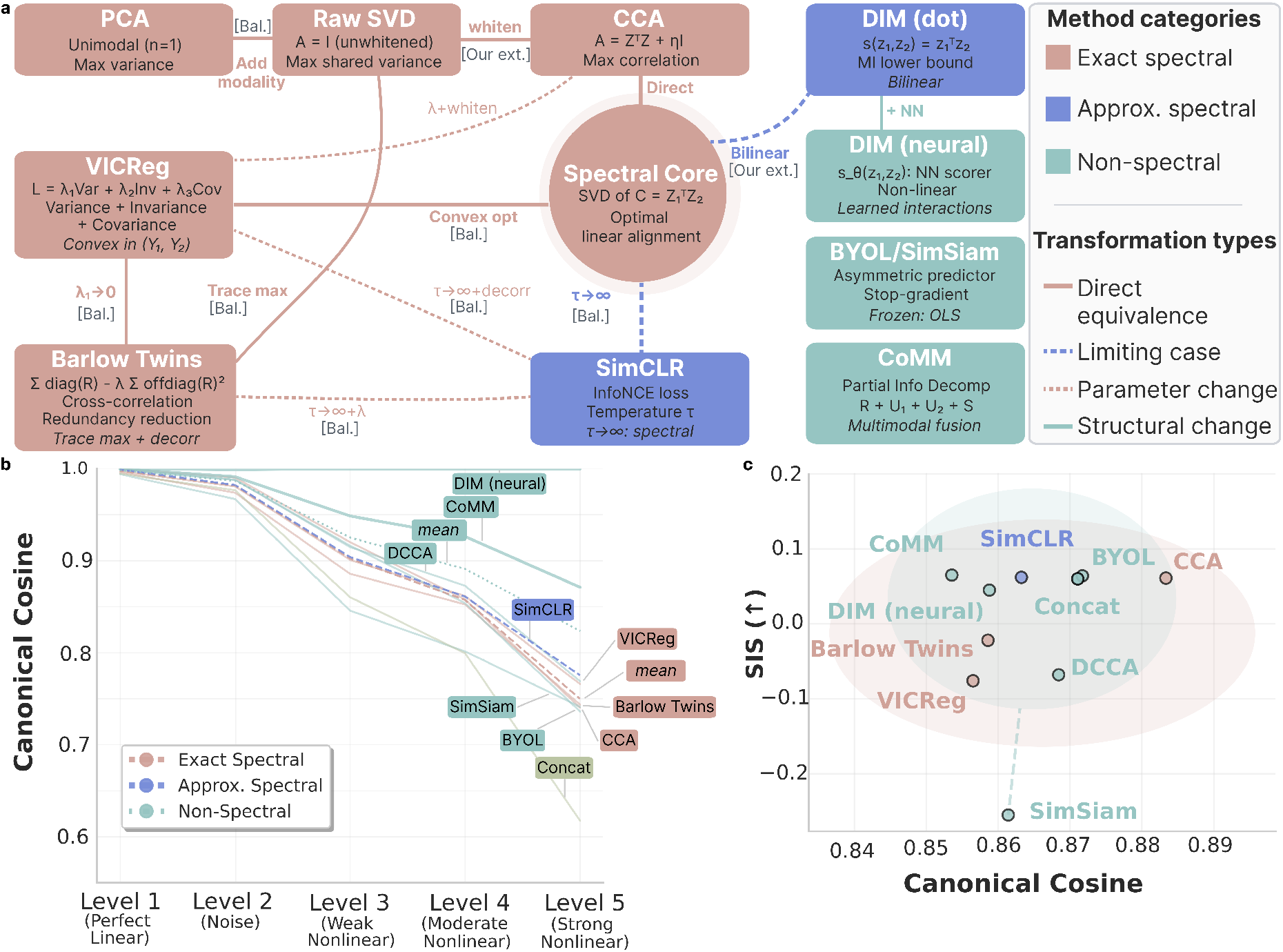
Assessing the spectral ceiling of the multimodal frozen encoder framework. **a**, Theoretical taxonomy of multimodal fusion objectives under frozen encoders. Exact spectral alignment methods (salmon) reduce to closed-form singular value decomposition/canonical correlation analysis (SVD/CCA) solutions of the cross-covariance matrix. Approximate spectral methods (purple) converge to the same solution in limiting regimes. Non-spectral integration methods (teal) introduce asymmetric predictors or synergy-aware terms that break the eigenvalue structure and permit nonlinear interactions. [Bal.] denotes connections made in Balestriero and LeCun [27] and [Our ext.] denotes connections made in our extension of their analysis. **b**, Synthetic evaluation under increasing nonlinearity (Methods). Canonical cosine measures similarity between the recovered shared signal and the ground-truth shared structure in the generator (higher indicates closer recovery). Solid lines show individual methods, dashed lines show category means, and shaded regions represent 95% confidence intervals. **c**, Alignment–synergy landscape on the lung dataset. Each point is a fusion method, plotted by the canonical cosine to the spectral (linear) solution (x-axis; higher means closer agreement with the SVD solution) and SIS (y-axis). This plot illustrates the empirical relationship between the proximity to the linear alignment solution and the measured multimodal gain under SIS for this dataset.

#### Non-spectral methods are robust to nonlinearity

We validate this distinction using synthetic data with controlled nonlinear coupling between modalities (Methods). In this simulation study, canonical cosine is computed against the generator’s ground-truth shared signal and therefore measures recovery of the underlying cross-modal relationship (Fig. 3b). While all methods recover the signal in linear regimes, spectral methods degrade rapidly as nonlinearity increases; whereas non-spectral methods degrade at a slower rate, maintaining substantially higher canonical cosine under strong nonlinearity. This confirms that spectral alignment is inherently limited to linear structure, while non-spectral objectives can capture nonlinear cross-modal interactions.

In real biological data, ground truth is unavailable. Here, canonical cosine instead quantifies similarity to the spectral reference solution (i.e., singular value decomposition/canonical correlation analysis; Fig. 3c). Under this interpretation, proximity to the spectral solution reflects recovery of *linear redundancy*, while deviations indicate representations that move beyond purely linear alignment. Consistent with the synthetic results, methods that remain close to the spectral solution do not necessarily achieve high SIS, whereas several non-spectral methods achieve higher SIS despite deviating from the linear optimum (Supplementary Fig. 5). Non-spectral methods deviate from the linear optimum to capture complementary structure, achieving higher SIS across datasets (Supplementary Fig. 5 and Supplementary Note 8).

### Scaling analysis reveals that unimodal fine-tuning is more sample-efficient than alignment

A central practical question for deploying cell foundation models is how to allocate limited labeled data: should one invest in adapting a strong unimodal expert, or in fusing multiple modalities through a multimodal interface? This tradeoff is especially relevant given the distinct information regimes identified in Fig. 1e, which distinguish tasks that are unimodal-sufficient from those that are genuinely crossmodal dependent. To resolve this empirically, we perform the following scaling analysis. First, we progressively fine-tune the gene expression encoder on increasing fractions of downstream data (1% to 100%), while keeping the image encoder frozen, thereby creating a controlled setting to compare unimodal adaptation against frozen multimodal fusion. We then evaluate whether multimodal fusion provides additive value over unimodal adaptation (Fig. 4 and Supplementary Note 9). This analysis directly investigates whether performance gains arise primarily from strengthening the dominant modality or from integrating complementary signal across modalities.

**Figure 4.**
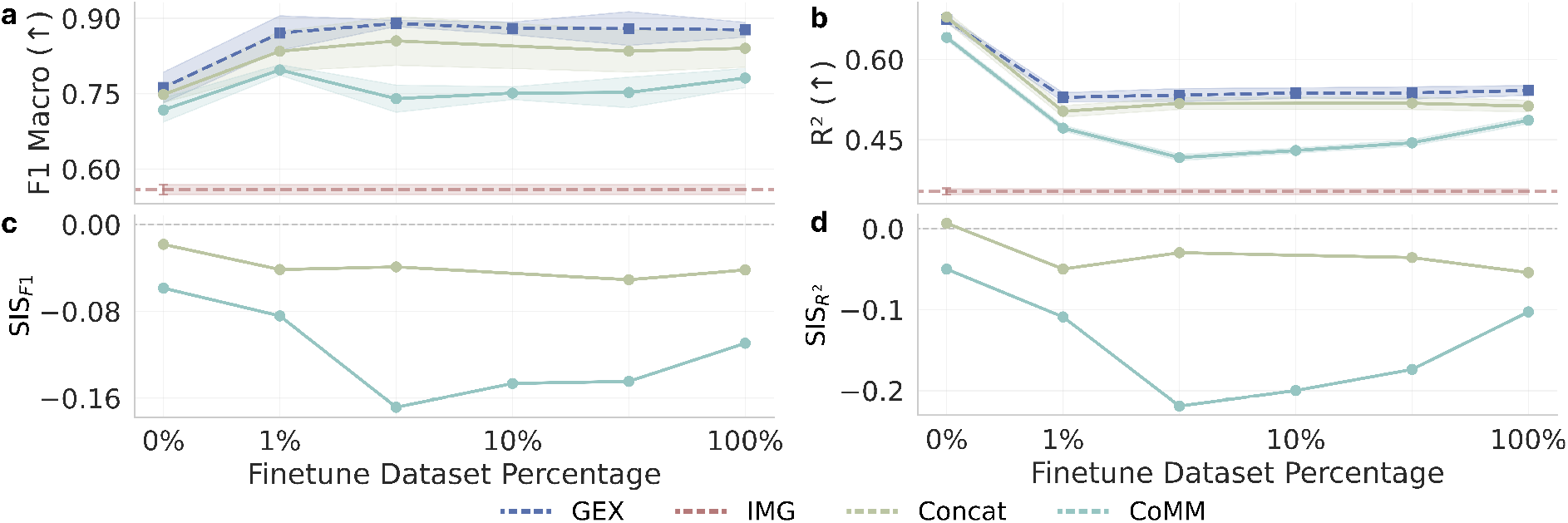
Scaling behavior of unimodal fine-tuning and multimodal fusion. **a-b**, Performance on the lung dataset as a function of labeled downstream training data used to fine-tune the GEX encoder (x-axis; depicted as log-scaled fractions). We report (a) F1 Macro for niche classification and (b) *R*^2^ for cell-type composition regression. Here, 0% indicates frozen encoders with only a linear probe trained, and *x*% indicates fine-tuning GEX on that fraction while keeping IMG frozen. **c-d**, Corresponding SIS values computed from the same probe metrics with panel (c) showing SIS_F1_ and panel (d) showing 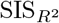. Curves show unimodal GEX and IMG baselines and two fusion interfaces (Concat and CoMM). Shaded regions denote the mean ± one standard deviation across five cross-validation folds.

#### Unimodal fine-tuning is highly sample-efficient

In the lung niche classification task, fine-tuning the GEX encoder yields rapid gains at small data fractions (Fig. 4a), with similar qualitative trends also seen in the breast and thymus datasets (Supplementary Fig. 6). Moving from frozen features (0%, where only the downstream probe is trained) to 1% fine-tuning substantially improves F1 Macro, after which performance largely saturates with additional data (Fig. 4a). Across all training fractions, the fine-tuned GEX encoder matches or exceeds the multimodal baselines (Concatenation and CoMM), indicating that adapting the strongest modality captures most attainable performance. A similar pattern holds for celltype composition regression, where the fine-tuned GEX encoder remains the strongest performer, while multimodal fusion yields only modest improvements relative to simple aggregation and does not close the gap to the unimodal baseline (Fig. 4b). Overall, these scaling curves indicate that, for this dataset and set of tasks, most attainable performance is captured by adapting the strongest unimodal expert, consistent with a unimodal-sufficient regime (Fig. 1e).

#### Fine-tuning reduces the incremental value of fusion under SIS

We next track SIS as the GEX expert is adapted (Fig. 4c,d). Once GEX is fine-tuned (from 1% onward), SIS is consistently negative for both tasks, indicating that fused representations do not outperform the best unimodal representation under the linear-probe evaluation used to define SIS. This behavior reflects a redundancy-dominated regime in which unimodal adaptation increasingly captures the accessible task-relevant signal, leaving little (linearly accessible) complementary information to be unlocked through integration. This indicates that, when the downstream task is dominated by information already present in a single modality, fine-tuning that expert is the most sample-efficient strategy. In such unimodal-sufficient regimes (Fig. 1e), frozen multimodal alignment introduces additional parameters without unlocking new task-relevant information.

#### Resolution mismatch preserves the value of integration

The scaling behavior on the thymus and breast datasets follows the information regimes identified in earlier sections (Supplementary Fig. 6 and Supplementary Note 10). In the thymus dataset, where gene expression is measured at coarse Visium resolution and modalities provide complementary information, multimodal integration remains competitive across all fine-tuning fractions. Although unimodal GEX performance improves gradually with fine-tuning, it does not clearly subsume the multimodal models, and the synergy-aware CoMM interface maintains stable performance and consistently positive SIS throughout. This identifies the thymus task as cross-modal-dependent, demonstrating that SIS correctly diagnoses a regime in which integration methods provide additive value beyond unimodal adaptation. Accordingly, under resolution mismatch and spatial ambiguity, complementary information from a second modality remains accessible even as the dominant expert is refined. In contrast, the breast dataset exhibits behavior more similar to the lung dataset, with high baseline performance and limited gains from doing either fine-tuning or fusion, indicating restricted space for additional improvement.

#### Scaling clarifies efficiency regimes for multimodal foundation models

Together, these results establish that SIS provides a principled diagnostic for determining whether unimodal adaptation suffices or if multimodal integration provides additive value. In the lung dataset (and similarly in breast), fine-tuning GEX rapidly saturates performance and drives SIS to become negative, indicating limited multimodal benefit under this setup and therefore a unimodal-sufficient regime (Fig. 4; Supplementary Fig. 6a-d). In the thymus dataset, where resolution mismatch induces cross-modal dependence, CoMM maintains positive SIS across training fractions, indicating a cross-modal-dependent regime (Supplementary Fig. 6e-h). In other words, when tasks are unimodal-sufficient, adapting the dominant expert provides the most sample-efficient path to optimal performance; however, when tasks are cross-modal-dependent, multimodal integration can expose complementary information that remains inaccessible to unimodal refinement alone. Thus, multimodal integration is not a universal performance amplifier, but a mechanism whose benefit depends on whether the downstream task fundamentally requires combining information distributed across modalities.

## Discussion

The ambition of constructing a “virtual cell”, a computational model that can predict cellular function across scales, requires more than aggregating multimodal measurements. It requires biological synthesis: combining complementary views to resolve ambiguity that cannot be resolved within any single modality. Yet, as unimodal datasets continue to grow and multimodal acquisition efforts begin to expand, current evaluation paradigms often conflate improvements from recovering redundant cross-modal structure with genuine integration. This conflation creates a practical blind spot: simple fusion objectives may appear effective even when they primarily propagate the strongest unimodal signal. To expose and investigate this conflation, we introduce the Synergistic Information Score (SIS) as a diagnostic for multimodal learning. By formalizing the distinction between alignment (recovering shared structure) and integration (making complementary information jointly accessible), SIS reveals that strong benchmark performance can arise even when a model effectively relies on a single dominant modality.

Our theoretical analysis identifies a mechanism that explains this pattern in a widely used practical workflow, where frozen backbones with a learned fusion interface are evaluated by linear probes. In this setting, frozen encoders fix the geometry of each modality, and many standard fusion objectives (including CCA-like or contrastive formulations under variance/whitening constraints) reduce to maximizing linear cross-covariance. This “spectral ceiling” is well suited to retrieval-focused objectives where modalities are expected to match in a shared latent space, but it also implies that fusion interfaces are biased toward extracting the linear redundancy already present in pretrained experts. As a consequence, these interfaces reliably recover shared signals while offering limited capacity to expose synergistic interactions that are not linearly expressible in the aligned features. Importantly, this ceiling is specific to the frozen encoder regime and to the common practice of assessing cross-modal alignment primarily with linear metrics. In end-to-end CLIP-style training, deep encoders can map complex correspondences into representations that appear approximately linearly aligned, although nonlinear modality gaps can persist in practice. Together, these considerations motivate objectives that go beyond geometric alignment and explicitly target information gain, such as CoMM.

Empirically, the alignment versus integration distinction translates into a concrete design principle for building virtual-cell-like models. In redundancy-dominated regimes with high cellular resolution and precise cross-modal correspondence (e.g., Xenium-like spatial profiling), unimodal experts already capture most task-relevant signal; correspondingly, SIS is near zero or negative. Here, our scaling analysis shows that unimodal fine-tuning is the most sample-efficient path to strong performance, and multimodal fusion provides little consistent improvement. In contrast, the promise of multimodal modeling lies in regimes where ambiguity or resolution mismatch distributes task-relevant information across modalities. For example, in spot-based spatial transcriptomics, coarse gene expression measurements and fine-grained histology create an integration regime in which simple aggregation can fail to make complementary signal accessible; similarly, predicting tissue context at an increasing spatial range shifts tasks away from local redundancy toward cross-modal dependence. In these settings, alignment alone is insufficient, and synergy-aware integration becomes competitive. Methods such as CoMM model cross-modal interactions rather than only enforcing agreement, enabling access to structure that is not recoverable from either modality alone under a linear probe.

These findings establish two information regimes that govern multimodal learning (Fig. 1e and Box 1), with SIS serving as a diagnostic to distinguish between them. In unimodal-sufficient regimes (SIS ≲ 0), task-relevant signal is already accessible within a single modality, and improving the strongest unimodal foundation model provides the most efficient path to performance gains. In cross-modal-dependent regimes (SIS > 0), complementary information becomes accessible through fusion, and multimodal fusion interfaces that model cross-modal interactions provide additive value. SIS therefore provides a practical diagnostic for guiding multimodal modeling: it identifies when unimodal refinement suffices and when integration is necessary to access complementary biological signal. This distinction is particularly relevant for synthesis-oriented settings, such as virtual cell efforts. Here, the objective is to combine information across biological scales rather than to recover redundant structure, aiming for regimes where fusion yields positive SIS (i.e., where cross-modal interactions become measurably accessible).

Ultimately, our work argues for a shift in multimodal learning from enforcing correspondence to achieving biological synthesis. A viable virtual cell model cannot be built by merely forcing modalities to correlate; it requires objectives that preserve each modality’s unique structure while making emergent properties accessible through interaction. We introduce SIS as a diagnostic probe for information regimes in multimodal learning and demonstrate that synergy-aware, integration objectives such as CoMM can outperform alignment objectives in cross-modal-dependent settings. Together, this contribution provides practical guidance for when to prioritize unimodal refinement versus multimodal integration in virtualcell-style modeling.

## Methods

### The Synergistic Information Score

In this section, we describe the overall problem statement that motivates the Synergistic Information Score (SIS). Given paired raw input data matrices from two modalities 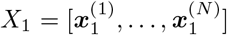 and 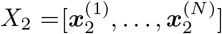 (e.g., histology images and gene expression profiles), together with a downstream task measured by the output vector 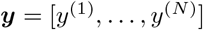 (e.g., niche labels or cell-type proportions), we define frozen unimodal representations as the following

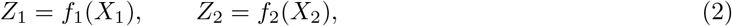

where *f*_1_ and *f*_2_ are frozen unimodal encoders and 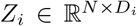 with *D*_*i*_ being the dimension of the embedded space. We next introduce the concept of a *fusion interface*, denoted by 𝒢= (*g*_1_, *g*_2_), which operates on the frozen unimodal representations. Here, 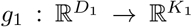 and 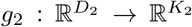 are modality-specific fusion mappings (also referred to as projection heads), yielding

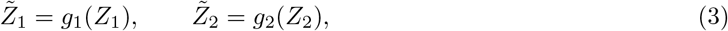

where 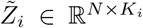 with *K*_*i*_ denoting the dimension of the modality-specific fusion. Depending on the objective, the fusion interface may instantiate an alignment-based or an integration-based fusion strategy. All fusion objectives considered in this work learn projection mappings *g*_1_ and *g*_2_ from unimodal representations (*Z*_1_, *Z*_2_), with optimization objectives defined on the resulting projected representations 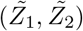. In the spectral analysis, we explicitly restrict to the *linear regime* where *g*_*i*_(*Z*_*i*_) = *Z*_*i*_*W*_*i*_, which enables closed-form eigenvector or singular value decompositions (SVD). Outside this regime, *g*_*i*_ may be nonlinear, but the spectral characterizations no longer apply. We then form a single fused representation

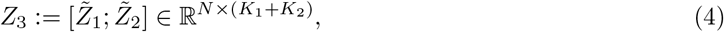

and train a linear probe *h* to predict the output vector ***y*** for downstream evaluations. As part of our motivation for SIS, the goal is to develop a scalar measure that has the following properties:

- It quantifies the *complementarity* between *Z*_1_ and *Z*_2_ with respect to a downstream task;
- It is *grounded in information theory* yet tractable in high-dimensional representation spaces;
- It enables *task-dependent comparison* across multimodal fusion methods. Below, we review key information-theoretic preliminaries before deriving the SIS.

#### Mutual information (MI)

The mutual information between a target random variable *Y* ∈ 𝒴 (where the output vector ***y*** contains realizations of *Y*) and a source random variable *Z* ∈ 𝒵 is defined as

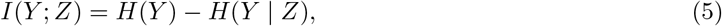

where *H*(*Y*) = − ∑_*y*_ *p*(*y*)log*p*(*y*) denotes the entropy of *Y* and *H*(*Y* | *Z*) denotes the conditional entropy. By the data processing inequality [41], for any (possibly randomized) decoding function 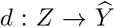, we have *I*(*Y* ; *d*(*Z*)) ≤ *I*(*Y* ; *Z*).

#### Partial information decomposition (PID)

Partial information decomposition [25, 26] decomposes the mutual information between a target variable *Y* and two source variables into redundant, unique, and synergistic components. In our setting, the sources are the projected representations 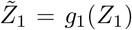 and 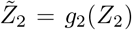 produced by the fusion interface, and we define the joint representation as their concatenation 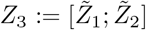. Since concatenation is a bijective reparameterization of the pair 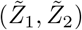, mutual information is invariant under this transformation, and we have

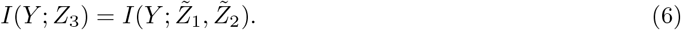

PID therefore yields

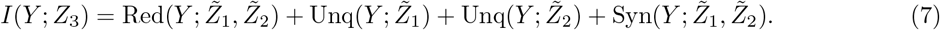

Here, the redundant component Red(Y ; 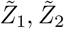) quantifies information in *Y* that can be explained by either projected representation alone, the unique components Unq(*Y* ; 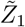) and Unq(*Y* ; 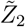) quantify information accessible exclusively from one projected representation, and the synergistic component Syn(*Y* ; 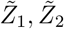) quantifies information in *Y* that is only accessible when both projected representations are considered jointly.

#### Deriving the Synergistic Information Score (SIS)

From Eq. (7) and by definition of PID, the mutual information between *Y* and a single source consists of the information it shares with the other source (redundancy) and the information that is unique to itself [25, 26]. Equivalently, synergy contributes only to the joint information *I*(*Y* ; 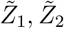) and not to *I*(*Y* ; 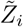) for either source individually:

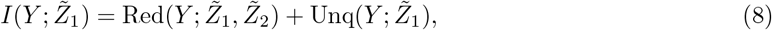

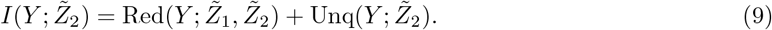

Solving for the synergistic component then yields

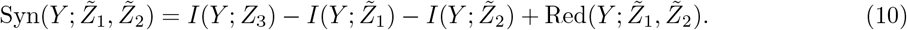

Following standard practice in the PID literature [25, 26], we approximate redundancy using the minimum mutual information (MMI) principle,

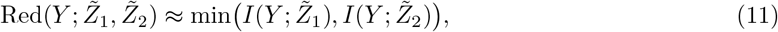

which yields the approximation

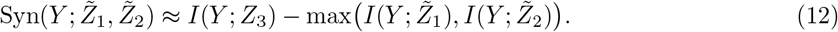

In practice, we compare fusion against the strongest *original* unimodal representation from either *Z*_1_ or *Z*_2_. Since each aligned representation 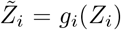 is a deterministic function of *Z*_*i*_, the data processing inequality implies 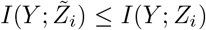 for *i* ∈ {1, 2}. Replacing max 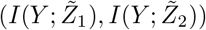 by the stronger baseline max(*I*(*Y* ; *Z*_1_), *I*(*Y* ; *Z*_2_)) therefore yields a conservative estimate of the fusion gain. We then define the *Synergistic Information Score (SIS)* as

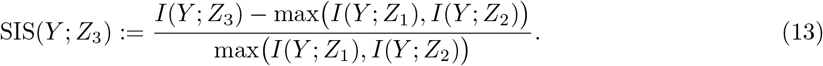

Intuitively, SIS measures the relative amount of task-relevant information made accessible by fusion beyond the strongest unimodal baseline under the chosen probe class. Positive SIS indicates accessible synergy, whereas near-zero or negative values indicate redundancy dominance or fusion degradation, respectively.

#### Note on approximation error

The MMI assumption is exact if the information in the “weaker” modality is a subset of the “stronger” modality (i.e., *Z*_min_ ⊆ *Z*_max_). The approximation error is given by Error = Unq(*Y* ; *Z*_min_). In our biological setting, where modalities are observed over the same structure (e.g., imaging and gene expression are taken from the same tissue), we assume substantial redundancy.

#### Boundedness

Here, we derive the range of SIS. Let *I*_max_ = max(*I*(*Y* ; *Z*_1_), *I*(*Y* ; *Z*_2_)). Using the subadditivity of mutual information, *I*(*Y* ; *Z*_3_) ≤ *I*(*Y* ; *Z*_1_) + *I*(*Y* ; *Z*_2_), we can specify the upper bound

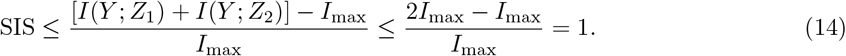

Since *I*(*Y* ; *Z*_3_) ≥ 0, the lower bound is SIS ≥ (0 − *I*_max_)*/I*_max_ = − 1. Thus, SIS ∈ [− 1, 1], where positive values indicate greater synergy and negative values indicate fusion failure (i.e., performance degradation).

#### Mathematical properties and important edge cases of SIS

The SIS satisfies the following mathematical properties:

- **Symmetry**. SIS(*Y* ; *Z*_3_) = SIS(*Y* ; 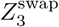), where *Z*_3_ := [*g*_1_(*Z*_1_); *g*_2_(*Z*_2_)] and 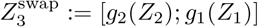. This holds because 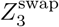 differs from *Z*_3_ only by a permutation of feature coordinates, and the linear probe hypothesis class used to estimate SIS is invariant under such permutations.
- **Sign**. SIS > 0 indicates synergy, while SIS *<* 0 indicates fusion degradation.
- **Boundedness:** SIS ∈ [− 1, 1] where *I*(*Y* ; *Z*_3_) ∈ [0, *I*(*Y* ; *Z*_1_) + *I*(*Y* ; *Z*_2_)] (see derivation above).
- **Task-specificity**. SIS depends on the target label *Y*, distinguishing tasks that truly require crossmodal reasoning.

In addition to these properties, there are a few edge cases that are worth noting:

- **Redundant modalities**. If *I*(*Y* ; *Z*_1_) = *I*(*Y* ; *Z*_2_) = *I*(*Y* ; *Z*_3_), then SIS = 0.
- **High synergy**. If *I*(*Y* ; *Z*_3_) = *I*(*Y* ; *Z*_1_) + *I*(*Y* ; *Z*_2_) (i.e., maximum possible synergy), then SIS approaches its upper bound of 1 (achieved exactly when *I*(*Y* ; *Z*_1_) = *I*(*Y* ; *Z*_2_)).
- **Fusion failure**. If *I*(*Y* ; *Z*_3_) *<* max(*I*(*Y* ; *Z*_1_), *I*(*Y* ; *Z*_2_)), then SIS *<* 0, indicating that fusion degrades performance below a better unimodal baseline.

#### Computationally efficient estimation of SIS via linear probes

For high-dimensional representations, computing the exact mutual information between the task variable *Y* and representation variable *Z* is intractable. We therefore approximate mutual information using the empirical performance of a linear probe *h* trained on a given representation matrix *Z*, denoted as Perf(*Z*). In our setting, *Z* is instantiated either as a unimodal representation (*Z*_1_ or *Z*_2_) or as the multimodal representation 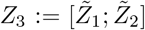. We then compute SIS as the following

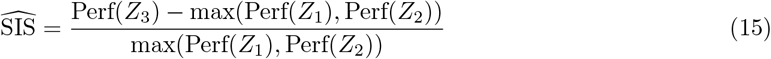

where performance is measured via a F1 Macro score for classification tasks, and performance is determined via R-squared (*R*^2^) for regression-based tasks. While standard performance metrics (like F1 scores) are not strictly additive in the same way as entropy, 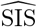 serves as a practical estimator of the *relative predictive gain* for several reasons:

- By the data processing inequality, any decoder 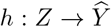 can only extract information present in the representation, meaning, *I*(*Y* ; *h*(*Z*)) ≤ *I*(*Y* ; *Z*) [41]. In our setting, *h* is a linear probe (classifier or regressor), so its performance reflects a lower bound on the task-relevant information in *Z* that is *linearly accessible* (also see discussion around Eq. (5)).
- In the context of foundation models, we care about *accessible* information that downstream tasks can readily extract, not just from an information-theoretic capacity.
- By restricting the probe to be linear, 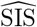 measures whether the fusion interface 𝒢 = (*g*_1_, *g*_2_) has rendered synergistic information *linearly accessible*, which is a useful metric for evaluating whether a foundation model provides ready-to-use representations. If synergy exists but is confined to a complex nonlinear manifold that a linear probe cannot access, then the interface has failed in its purpose.

Therefore, our proposed approach provides a conservative estimate of the amount of task-relevant synergistic information that is linearly accessible from the fused representation. If a linear probe cannot extract information, the fusion has failed to make synergistic patterns readily usable. Again, a positive SIS indicates the extraction of accessible synergistic states; whereas a near-zero or negative SIS implies redundancy or fusion failure, respectively. In the main text, we denote the empirical estimator 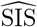 simply as SIS when no ambiguity arises, and specify the probe-dependent metric via indices such as SIS_F1_ and 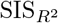.

### Spectral alignment theory

We extend the spectral theory of self-supervised learning to the multimodal setting with frozen encoders. Balestriero and LeCun [27] showed that diverse unimodal self-supervised learning (SSL) objectives (e.g., VICReg, Barlow Twins, SimCLR) admit spectral interpretations and derived closed-form optima in the linear regime under standard variance/whitening constraints. HaoChen et al. [23] further analyzed a *spectral contrastive loss* and connected it to contrastive learning objectives from the augmentation graph perspective. These works motivate that, when representations are fixed, broad classes of SSL objectives reduce to constrained spectral optimization problems. As part of our contributions, we adapt this viewpoint to analyze *multimodal alignment* under frozen encoders, where the goal is to couple representations from two different fixed-modality encoders rather than augmented views of the same input.

#### Spectral problem setup

We adopt the same notation and fusion interface defined earlier in this section. Frozen encoders *f*_1_ and *f*_2_ produce representations 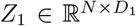 and 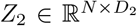. A fusion interface 𝒢 = (*g*_1_, *g*_2_) maps these to projected representations 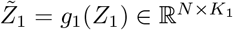 and 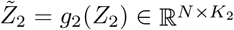. Here, we analyze fusion objectives that are defined on the paired projected representations 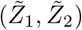.

#### Spectral methods primer

Spectral methods analyze matrices through their eigenvalues and eigenvectors, which identify orthogonal directions capturing dominant variance and correlation structure (see Chung [42] for a textbook summary). For paired representations 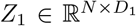 and 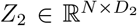, their shared structure is characterized by the cross-covariance 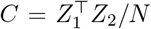. The SVD of this matrix *C* = *U* Σ*V* ^⊤^, yields orthogonal directions that maximize alignment between the two spaces, with singular values quantifying alignment strength. These directions arise as solutions to constrained alignment problems of the form 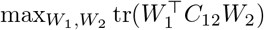 subject to orthonormality constraints, which are solved by the top singular vectors (Eckart-Young theorem [40]). Whitening or variance constraints ensure nondegenerate solutions and isolate correlation structure. Thus, under linear projections and variance normalization, alignment objectives reduce to identifying the dominant singular vectors of cross-covariance matrices, which define the optimal shared subspace between representations.

#### The linear regime

To obtain closed-form spectral characterizations, we restrict to the *linear regime* in which the fusion mappings are linear maps applied row-wise,

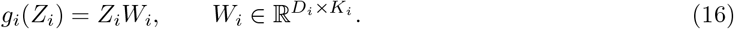

This restriction is used solely for theoretical purposes. Note that in the general setting, *g*_*i*_ may be nonlinear, in which case the spectral closed-form solutions derived below no longer apply. We do not assume that the relationship between modalities is linearly realizable in the frozen feature space; rather, the linear-regime analysis characterizes the optimizer of a broad class of alignment objectives under linear heads. Moreover, variance/whitening (or analogous) constraints are essential to exclude collapsed solutions (e.g., *W*_1_ = *W*_2_ = 0) and yield non-trivial spectral structure. This distinction matters because:

- **Frozen geometry**. Unlike end-to-end training, frozen encoders impose a fixed representation geometry.
- **Constrained spectral optima**. Under linear heads and standard variance/whitening constraints, several covariance-based objectives (e.g., VICReg, Barlow Twins) admit spectral solutions [27]. Closely related spectral perspectives for contrastive objectives have also been studied [23].

#### The spectral ceiling

We define the *spectral ceiling* as the constrained optimum attained by redundancy-maximizing alignment objectives in the linear-head and frozen encoder regime. While practical implementations may use nonlinear heads, the linear derivation below establishes a reference bound for objectives whose solutions concentrate on cross-covariance structure. In the linear regime, a canonical constrained trace maximization problem is

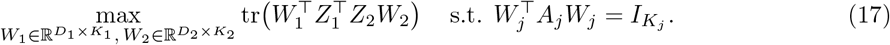

This problem is solved by the singular value decomposition (SVD) of an appropriately whitened cross-covariance matrix (via the Eckart–Young theorem [40]). Here, 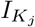 denotes the *K*_*j*_ × *K*_*j*_ identity matrix. For our frozen encoder setting, we state this classical result:

##### Proposition 1

(Spectral solution for frozen encoder alignment). *Assume Z*_1_, *Z*_2_ *are column-centered. The trace maximization problem in* Eq. (17) *with constraints* 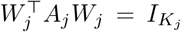, *where K*_*j*_ *is the j-th target dimension and A*_*j*_ ⪰ 0, *is solved by the top K* = min(*K*_1_, *K*_2_) *singular vectors of the whitened cross-covariance*

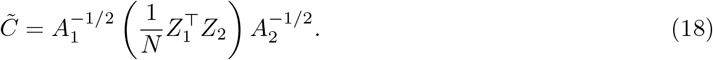

*Let* 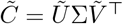 *be the SVD. Then one optimal choice is*

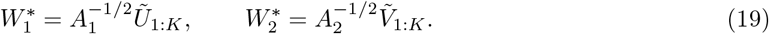

*Proof*. This follows directly from the Eckart–Young theorem [40]. The canonical correlation analysis (CCA) case corresponds to 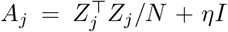 as established by Hotelling [43], with *η* denoting a regularization hyperparameter.

#### Categorization of fusion methods

In the main text, we categorize methods by whether their objectives reduce to Eq. (17) in the linear limit (also see Fig. 3a), following the spectral viewpoints in Balestriero and LeCun [27] and related analyses of contrastive objectives in HaoChen et al. [23]. We refer to methods whose linear-limit objectives admit closed-form spectral solutions as *spectral methods*. We then further classify approaches whose linear-limit objectives reduce directly to the closed-form spectral solutions under standard variance/whitening (anti-collapse) constraints as *exact spectral methods*.

- **Linear CCA** [43]: Directly solves a generalized eigenvalue problem.
- **VICReg** [35] and **Barlow Twins** [44]: With frozen representations and linear heads, these objectives admit constrained spectral solutions [27].

We describe *approximately spectral* methods as objectives whose optimization is closely tied to spectral structure in certain limiting regimes (e.g., large-batch, many-negative settings for contrastive learning), while we avoid requiring an exact reduction for the specific loss used in practice.

- **SimCLR (InfoNCE)** [36]: A contrastive objective with close connections to spectral viewpoints under appropriate limits and assumptions (see, e.g., HaoChen et al. [23] for a spectral contrastive formulation and related analysis).

Lastly, we describe *non-spectral* methods as objectives whose structure prevents reduction to a symmetric trace maximization eigenproblem. This class includes the following list.

- **Naive Concatenation (Concat)**: Joins representations without learning *g*_1_ and *g*_2_. Since it does not perform redundancy reduction or subspace selection, it preserves all information (redundant and unique) and does not collapse to the spectral subspace. In the main text, we also refer to Concat as Aggregation, as it aggregates information from both modalities.
- **Deep CCA (DCCA)**: Optimizes the canonical correlation objective using nonlinear projection heads. Unlike linear CCA, the nonlinear transformations allow it to capture correlations on a learned manifold, thereby escaping the linear cross-covariance spectrum of the frozen inputs.
- **DIM (Neural)** [45]: Maximizes mutual information using a parameterized neural critic (discriminator) rather than a simple dot-product. This neural estimation allows it to capture complex nonlinear dependencies that do not reduce to trace maximization.
- **BYOL** [46] and **SimSiam** [47]: Uses asymmetric prediction with stop-gradient. With frozen encoders, this becomes an asymmetric regression problem rather than a symmetric trace maximization.
- **CoMM** [34]: Uses a PID-aware objective with specific synergy terms that explicitly deviate from redundancy maximization.

Additional details for each method are provided in Supplementary Note 4.

#### Empirical analysis of spectral convergence

To empirically test our categorization, we quantify how closely each method’s learned projected embeddings 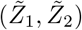 adhere to the linear spectral solution, even when using nonlinear projection heads. We compute a reference linear solution via the SVD of the centered cross-covariance 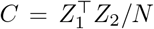 on *N* = 1000 paired patches from the held-out test set. Let *C* = *U* Σ*V* ^⊤^ and denote the top-*K* singular subspaces by *U*_1:*K*_ and *V*_1:*K*_. We use parallel analysis [48] to estimate an effective dimensionality *K* by permuting rows of *Z*_2_ independently *L* = 100 times to break pairing while preserving marginals, and taking the count of observed singular values exceeding the 95^th^ percentile of the null singular values. Note that, in our experiments, this yielded *K* = 5 for which we verified stability across random seeds and train/test splits.

#### Convergence metrics

We evaluate adherence to the reference spectral subspaces using: (i) a Procrustes-style subspace distance, (ii) projection energy onto the reference subspace, and (iii) canonical cosine or principal angle-based similarity. Exact definitions and implementation details are given in Supplementary Note 8. A higher canonical cosine indicates closer adherence to the linear redundancy solution, whereas a lower canonical cosine indicates deviation from this linear subspace (which can reflect either nonlinear structure capture or failure, depending on downstream SIS).

#### Synthetic data generator

We designed a 5-level synthetic data generator that progressively increases nonlinearity and noise, enabling controlled evaluation of spectral versus non-spectral methods. The rationale behind this design is to create a systematic progression from perfect linear alignment to strong nonlinear coupling, with each level characterized by intrinsic complexity measures rather than target canonical cosine values. All levels use a shared latent structure: we generate a shared latent 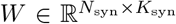 from a standard Gaussian, where *N*_syn_ = 20000 and *K*_syn_ = 10. Both modalities are constructed via orthonormal projection bases 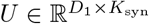 and 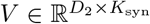 (with *D*_1_ = *D*_2_ = 128), scaled to achieve canonical correlations.

- **Level 1 (perfectly linear)**. Here, we generate a perfect linear relationship where *Z*_1_ = *WU* ^⊤^ and *Z*_2_ = *WV* ^⊤^, with no private noise (residual standard deviation *σ*_1_ = *σ*_2_ = 0). In other words, the signal-to-noise ratio is infinite. This provides an ideal baseline for spectral methods.
- **Level 2 (noise degradation)**. This scenario has the same linear structure as Level 1, but with increasing levels of private noise *σ*_1_, *σ*_2_ ∈ {0.0, 0.1, 0.2, 0.3, 0.5, 0.8, 1.0} added independently to each modality: *Z*_1_ = *WU* ^⊤^ + ***ε***_1_ and *Z*_2_ = *WV* ^⊤^ + ***ε***_2_ where 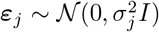. This creates a systematic signal-to-noise ratio degradation going from infinite to finite values.
- **Level 3 (linear with private structure)**. Here, we generate a linear relationship with structured private noise (where standard deviations are set to *σ*_1_ = *σ*_2_ = 0.3). This represents the case where modalities share a common signal, but each also contains modality-specific information (signal-to-noise ration ≈ 3.3).
- **Level 4 (moderate nonlinear coupling)**. In this case, we apply a nonlinear transformation to the second modality such that *Z*_1_ = *WU* ^⊤^ + ***ε***_1_ and *Z*_2_ = [tanh(*W*) + 0.2 sin(*W*)]*V* ^⊤^ + ***ε***_2_ with *σ*_1_ = *σ*_2_ = 0.3. The combination of tanh saturation and sinusoidal modulation introduces moderate nonlinearity (compositional structure: saturation ◦ periodic) while preserving some monotonicity.
- **Level 5 (strong nonlinear coupling with modality rotation)**. This last scenario has the strongest nonlinearity via a rotated nonlinear transformation to the second modality such that *Z*_1_ = *WU* ^⊤^ + ***ε***_1_ and *Z*_2_ = [tanh(*WR*)]*V* ^⊤^ + ***ε***_2_ where 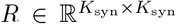 is a random rotation matrix. Here, we use increased asymmetric noise (*σ*_1_ = 0.5, *σ*_2_ = 0.7). This creates substantial nonlinearity (compositional structure: saturation ◦ rotation) and breaks linear alignment structure. The asymmetric noise (*σ*_1_ ≠ *σ*_2_) tests robustness to modality imbalance.

This progression systematically tests method robustness: spectral methods should excel at the first two levels but degrade with increasing nonlinearity, while non-spectral methods should maintain higher canonical cosine across all levels. The resulting canonical cosine values are measured empirically, not engineered to specific targets. We extend the spectral theory of self-supervised learning to the multimodal setting with frozen encoders. Balestriero and LeCun [27] showed that diverse unimodal self-supervised learning (SSL) objectives (e.g., VICReg, Barlow Twins, SimCLR) converge to spectral solutions recovering eigenmodes of kernel matrices. HaoChen et al. [23] further showed that the spectral contrastive loss concentrates on the top eigenfunctions of the data distribution. These works established that many seemingly different SSL objectives reduce to trace maximization problems when data representations are fixed. As part of our contributions in this work, we adapt this framework to analyze *multimodal alignment* under frozen encoders. This is a distinct setting from unimodal SSL, as it involves fusing representations across two different fixed-modality encoders rather than augmented views of the same input.

### Downstream task evaluation on real data

We evaluate unimodal and fused representations across five biologically grounded tasks with increasing spatial complexity. All downstream evaluations use linear probes trained on frozen representations.

- **Cell-type composition regression**. We predict the relative abundance of cell types in each patch using ridge regression. The target variable is a vector of cell-type proportions (excluding samples without annotations), and performance is reported via *R*^2^.
- **Niche type classification**. We classify patches into discrete tissue niche categories (e.g., tumor, stroma, immune) using multinomial logistic regression. Performance is reported via F1 Macro.
- **Niche consistency**. A pairwise consistency task testing whether patches sharing the same niche label are embedded closer than patches from different niches. Positive pairs are patches with identical niche labels, while negative pairs are patches with different niche labels. We train a logistic regression probe on absolute embedding differences to predict whether a pair shares the same niche, and report the area under the curve (AUC).
- **Spatial consistency**. A pairwise consistency task testing whether spatially nearby patches from the same slide are embedded closer than spatially distant patches. Positive pairs are patches within the *k*-nearest neighbors (default *k* = 5) on the same slide, while negative pairs are pairs beyond an 80th-percentile distance threshold on the same slide. We train a logistic regression probe on absolute embedding differences and report the AUC.
- **Spatial neighborhood prediction**. We predict properties of neighboring patches from center-patch embeddings across increasing distance bins. For classification, we predict the majority niche label among neighbors from the center embedding (via F1 Macro). For regression, we predict the mean cell-type composition of neighbors (via *R*^2^). Performance is reported across distance bins to quantify degradation with increasing spatial separation.

Throughout, we treat linear-probe performance as a practical proxy for task-relevant information accessible from a given frozen representation. Cross-validation or predefined train/validation/test splits are used as appropriate for each task.

### Model training and experimental details

Frozen embeddings are extracted from pretrained foundation models: UNI-200M (UNI-2) for histopathology images [5] and Nicheformer for spatial transcriptomics [8]. Patches are sampled from full slides and paired at the patch level. We filter samples with NaN embeddings and handle missing annotations according to each task definition.

#### Training of the fusion-interface

For each multimodal method, we train the fusion mappings (projection heads) *g*_1_ and *g*_2_ on the training split using PyTorch Lightning. Methods with self-supervised fusion objectives (e.g., CoMM, SimCLR, Barlow Twins, VICReg, BYOL, and SimSiam) use all paired samples in the training split (without downstream labels), while evaluation always uses held-out data. We use early stopping (patience 50 epochs) based on a smoothed validation loss. For linear CCA, we fit the closed-form solution on the training split. All learned methods are trained with AdamW (learning rate and weight decay selected via a sweep) and cosine annealing learning-rate schedules. See Supplementary Note 4 for more details.

#### Linear probe evaluation

After training the fusion interface, we train linear probes (logistic regression for classification; ridge regression for regression) on frozen representations (*Z*_1_, *Z*_2_, or *Z*_3_). We use 5-fold cross-validation or predefined train/validation/test splits depending on the evaluation setting. For spatial tasks, we enforce cross-slide validation to prevent spatial leakage across splits.

#### Spectral analysis

Reference spectral subspaces are computed from *N* = 1000 held-out test patches via the SVD of the centered cross-covariance 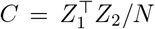. We use parallel analysis with *L* = 100 permutations to determine an effective dimensionality (typically *K* = 5). Canonical cosine between learned and reference subspaces is computed following standard numerical linear algebra protocols [49]; exact definitions are provided in Supplementary Note 8.

## Data availability

Datasets used in this study are publicly available from their original sources as detailed in the Supplementary Notes. Pretrained model resources used for feature extraction include the Nicheformer gene expression (GEX) encoder [8], trained on the publicly available SpatialCorpus-110M dataset, and the UNI-2 imaging (IMG) encoder [5], for which pretrained weights are publicly released while the underlying pretraining data are subject to institutional and regulatory constraints. Detailed descriptions of pretraining data sources and model checkpoints are provided in Supplementary Note 2.

## Code availability

The code for computing the Synergistic Information Score (SIS), spectral alignment analysis, and all multimodal alignment methods evaluated in this study is publicly available at https://github.com/microsoft/cell_synergy. The repository includes implementations for all benchmarked methods, preprocessing pipelines, and evaluation scripts to reproduce all results presented in this manuscript.

## Acknowledgments

We thank the Biomedical Machine Learning (BioML) group at Microsoft Research and members of the Theis Lab for thoughtful suggestions and useful conversations on previous drafts of the manuscript. We would like to thank Merel Kuijs, Rushin Gindra, Korbinian Träuble, and Christian Matek for their support with the multimodal datasets used in this study. We also thank Faisal Mahmood (Harvard) and his lab for assistance with the UNI-2 model. TR is supported by the Helmholtz Association through the Munich School for Data Science. TR was also supported by the German Federal Ministry of Research, Technology and Space (BMFTR) under grant no. #01IS18053A. FJT acknowledges support from the European Union (ERC, DeepCell – #101054957). SR acknowledges funding support from NCI K08 CA260442. Any opinions, findings, and conclusions or recommendations expressed in this material are those of the author(s) and do not necessarily reflect the views of any of the funders.

## Author contributions statement

TR and LC jointly designed the study and developed the theory and methods. TR and EZ implemented the method and performed the analyses. JH helped with developing theory. TR, EZ, and LC wrote the initial draft of the manuscript. FJT, SR, PSW, APA, and LC supervised the project. SR, PSW, APA, and LC provided resources. All authors contributed to interpreting the results and revising the manuscript.

## Supplementary Notes

### 1 Related work: histopathology-gene expression integration

Integrating histopathology with spatially resolved gene expression is an active area of research spanning (i) predictive models that infer expression (or cell states) from images, (ii) paired representation learning objectives for image-expression data, and (iii) downstream analysis pipelines that combine both modalities for specific biological questions and downstream tasks. In this section, we summarize representative directions to contextualize our benchmarking effort and clarify how our study differs.

#### Predicting spatial gene expression from histology

A major line of work trains models to reconstruct or predict spatial gene expression directly from Hematoxylin and Eosin (H&E) stains using paired patches/spots as supervision. Representative examples include Hist2ST [50], BLEEP [51], and DeepSpot [52]. Some related work also predicts bulk or slide-level expression from histology (e.g., HE2RNA [53]) while others synthesize histology conditioned on transcriptomic profiles (e.g., cascaded diffusion models [54]). These approaches primarily target *cross-modal prediction* (image → expression) rather than diagnosing *when* multimodal fusion increases task-relevant information beyond the strongest unimodal encoder.

#### Paired image-expression representation learning (fusion objectives)

Several works use paired histology and expression profiles to learn a joint embedding space via contrastive or correlation-based objectives. For example, TANGLE learns transcriptomics-guided slide representations via multimodal pretraining [55], and resources such as HEST-1k enable paired training and evaluation at scale [56].

In our framework, when such objectives are applied in the common *frozen encoder + learned interface* workflow, they are expected to preferentially recover redundant cross-modal structure, motivating our spectral analysis.

#### End-to-end multimodal models

Recent efforts aim to scale multimodal training across cohorts and tissue types. For example, H&Enium [57] and OmiCLIP [58] train end-to-end components on large paired pathology-transcriptomics resources. Because these models train multiple components jointly (encoders and fusion), it is challenging to attribute gains to a specific fusion interface and to disentangle redundancy recovery from synergistic integration without targeted ablations.

#### Analysis frameworks combining histology and spatial transcriptomics

A complementary ecosystem of methods integrates histology with spatial transcriptomics for specific downstream analyses (e.g., predicting tumor microenvironment structure or multi-resolution integration) like Starfysh [59], while other methods aim to perform platform-style integration via image registration such as VoltRon [60]. These tools are typically optimized for particular downstream analyses, rather than isolating and comparing fusion objectives under fixed unimodal backbones as done in our work here.

#### Connections to the broader multimodal alignment literature

Optimal transport approaches such as SCOT align modalities by matching relational structure via (Gromov-)Wasserstein objectives and are particularly useful when feature spaces differ and correspondences are not well captured by linear correlations [61]. Conceptually, these methods address the same challenge of correspondence under mismatch; however, they are developed for single-cell multi-omics alignment settings and do not target the frozen encoder to fusion-and-probe workflow evaluated in our benchmark. We therefore treat them as complementary approaches and do not include them in the empirical comparison.

#### Key distinctions of our work

Most prior work proposes complete models or task-specific pipelines. In contrast, we (i) hold strong pretrained unimodal encoders fixed, (ii) isolate the *fusion interface* as the unit of comparison, and (iii) evaluate across tasks that probe different information regimes (redundancy, modality-unique structure, and synergy). This yields actionable guidance: alignment-style objectives are appropriate when the goal is to recover shared structure, simple aggregation preserves modality-specific signals, and synergy-aware interfaces are needed when task-relevant information emerges only through cross-modal interaction.

#### Choice of unimodal encoders

Our experiments use UNI-2 [5] for histology and Nicheformer [8] for spatial gene expression. Other strong pathology encoders pretrained with contrastive or vision-language objectives (e.g., CONCH [62] or Virchow2 [7], as well as other pathology vision-language models) could be substituted on the imaging side, and alternative transcriptomic foundation models (e.g., scGPT [9], Geneformer [4], or UCE [63]) could be substituted on the gene expression side. Because our analysis fixes unimodal encoders and isolates the fusion objective and probe-accessible information, we expect the qualitative regimes identified by SIS (unimodal-sufficient versus cross-modal dependent) to be robust to the specific choice of unimodal backbone, although absolute task performance may shift.

### 2 Datasets

We evaluated all fusion methods on three human spatial transcriptomics datasets to validate robustness and assess how dataset characteristics influence the utility of multimodal integration. The dataset was obtained on 01 July 2026 from the publicly available HuggingFace repository [29, 31, 33]. Briefly, the authors integrated spatial transcriptomics measurements (10x Xenium and Visium) with matched histopathology image patches and curated expert-defined niche annotations across three tissues (lung, breast, and thymus). This yielded a partially paired multimodal dataset with aligned image and gene expression embeddings and tissue-specific ground-truth region labels. These datasets differ systematically in size, spatial resolution, data acquisition technology, and label availability (Supplementary Table 1), enabling us to test our theoretical predictions across distinct information regimes.

**Supplementary Table 1.**
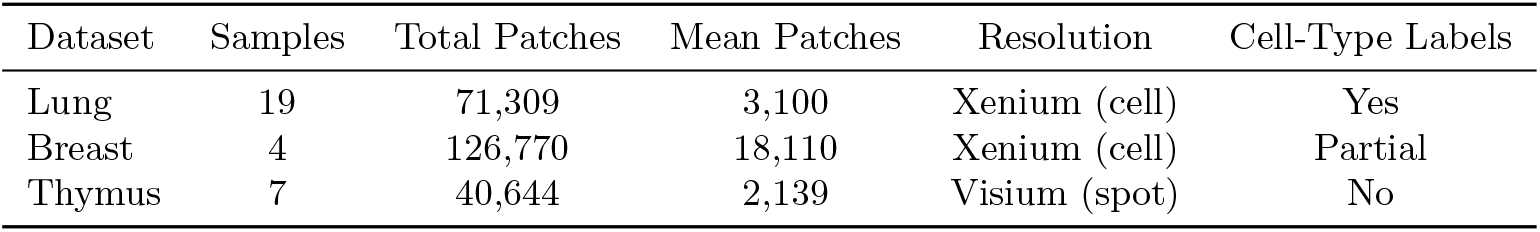
Dataset overview and general details.

| Dataset | Samples | Total Patches | Mean Patches | Resolution | Cell-Type Labels |
| --- | --- | --- | --- | --- | --- |
| Lung | 19 | 71,309 | 3,100 | Xenium (cell) | Yes |
| Breast | 4 | 126,770 | 18,110 | Xenium (cell) | Partial |
| Thymus | 7 | 40,644 | 2,139 | Visium (spot) | No |

#### Sample size and diversity

The lung dataset comprises 71,309 patches across 23 samples (mean 3,100 patches/sample) from a cohort of pulmonary fibrosis (PF) patients [28, 29]. The breast dataset contains 126,770 patches across 7 samples (mean 18,110 patches/sample) derived from patients with invasive ductal carcinoma (IDC) and invasive lobular carcinoma (ILC) [32, 33]. The thymus dataset contains 40,644 patches across 19 samples (mean 2,139 patches/sample) representing fetal and early postnatal development [30, 31]. As shown in Supplementary Figs. 1-3, the datasets exhibit distinct niche and cell type distributions that reflect their underlying biology.

#### Data acquisition and resolution

The lung and breast datasets utilize Xenium *in situ* technology, offering subcellular resolution (approximately 0.21 *µ*m/pixel) with precise spatial correspondence between histopathology and gene expression. In contrast, the thymus dataset uses Visium spatial transcriptomics, which aggregates expression within 55 *µ*m spots (capturing typically 1-10 cells per spot). This fundamental difference in spatial resolution creates distinct integration challenges: Xenium data presents a redundancy-dominated regime where morphology and expression are tightly aligned, while Visium data presents a resolution-mismatched regime requiring learned integration to bridge the scale gap between cellular histology and multi-cellular expression aggregates.

**Supplementary Figure 1.**
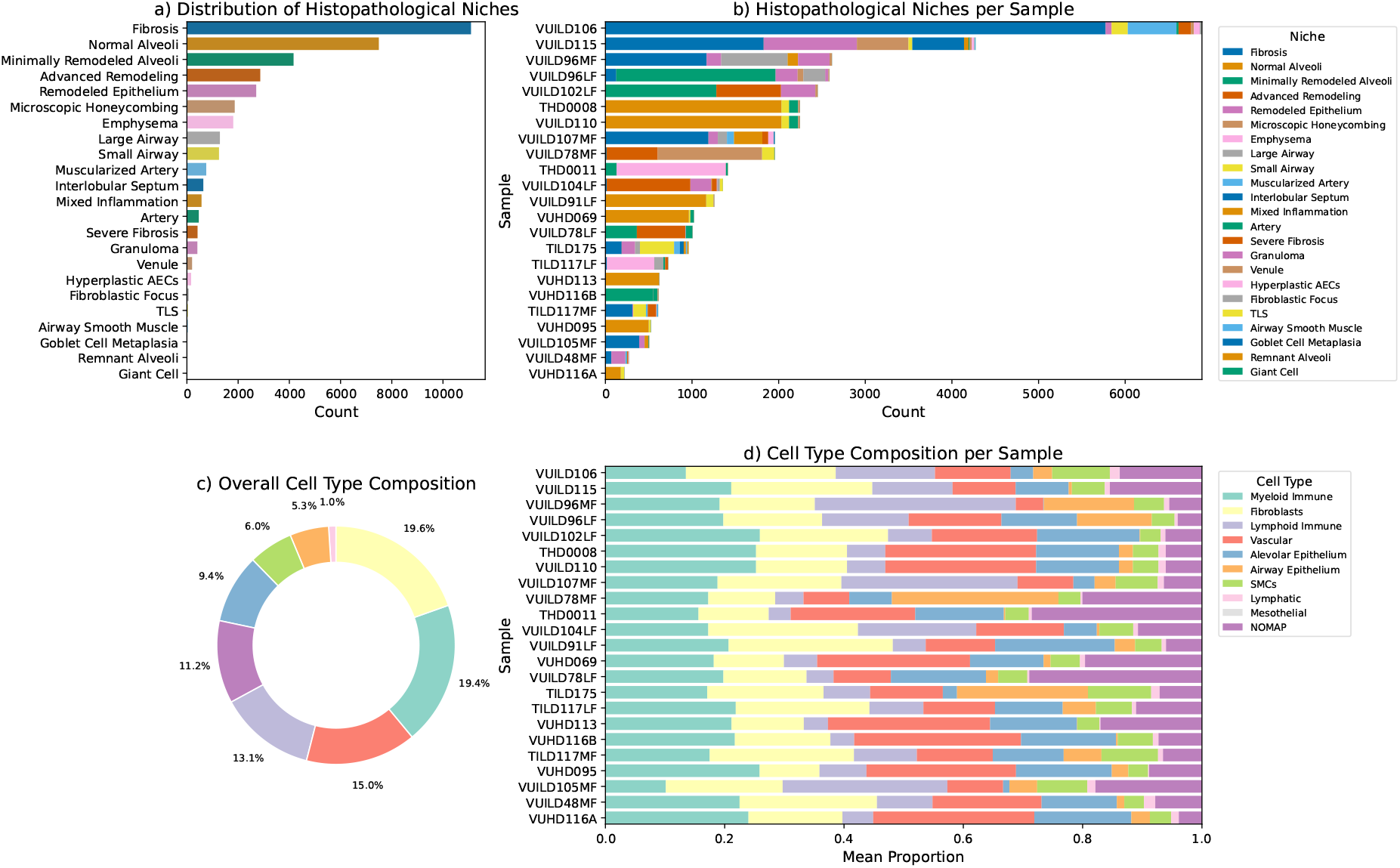
Data exploration for the lung dataset. **(a)** Overall distribution of histopathological niches, excluding *NOANNOT* patches without niche annotation. The lung dataset exhibits a characteristic fibrotic pattern with structural niches (alveolar regions, bronchioles) and pathological features (fibrosis, inflammation). **(b)** Niche distribution per sample shows heterogeneity across patients (2-13 distinct niches per sample). **(c)** Overall cell-type composition showing the dominance of structural cells (fibroblasts, epithelial), with immune infiltration (macrophages, lymphocytes). **(d)** Persample cell-type composition reveals patient-specific immune responses and fibrotic progression patterns.The *NOMAP* category represents cells excluded from evaluation.

#### Label availability and annotation

For the lung dataset, we utilized the 24 histological niche definitions provided by the original studies, which characterize distinct fibrotic regions including alveolar structures, bronchioles, vascular regions, and pathological features (fibrosis, inflammation, immune aggregates) [28, 29]. For the thymus dataset, we used the 11 morphological regions defined in the Human Thymus Cell Atlas, capturing developmental zones critical for T-cell maturation (cortex, medulla, corticomedullary junction, vasculature) [30, 31]. For the breast dataset, a collaborating pathologist annotated 14 distinct histological features across the samples based on morphological evidence and the Elston-Ellis scoring system for an unpublished study, including tumor grades, stromal patterns, immune infiltration, and microarchitectural features [33, 64]. As the thymus dataset lacks single-cell resolution and the breastdataset contains partially unlabeled regions, cell-type composition analysis was restricted to the lung dataset.

**Supplementary Figure 2.**
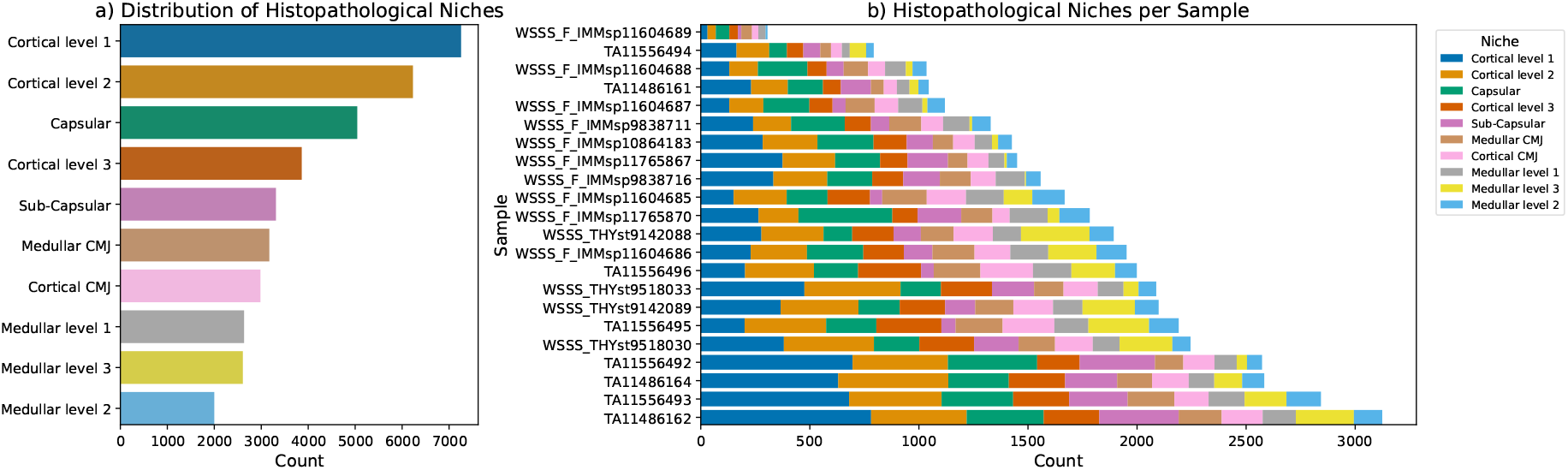
Data exploration for the thymus dataset. **(a)** Overall niche distribution excluding unassigned regions. The thymus exhibits highly structured developmental niches (cortex, medulla, cortico-medullary junction) reflecting T-cell maturation zones. **(b)** Niche distribution per sample demonstrates consistent developmental architecture across fetal and postnatal stages (10-11 distinct niches per sample, highest heterogeneity among all datasets). No cell-type composition labels are available for this spot-based Visium dataset.

**Supplementary Figure 3.**
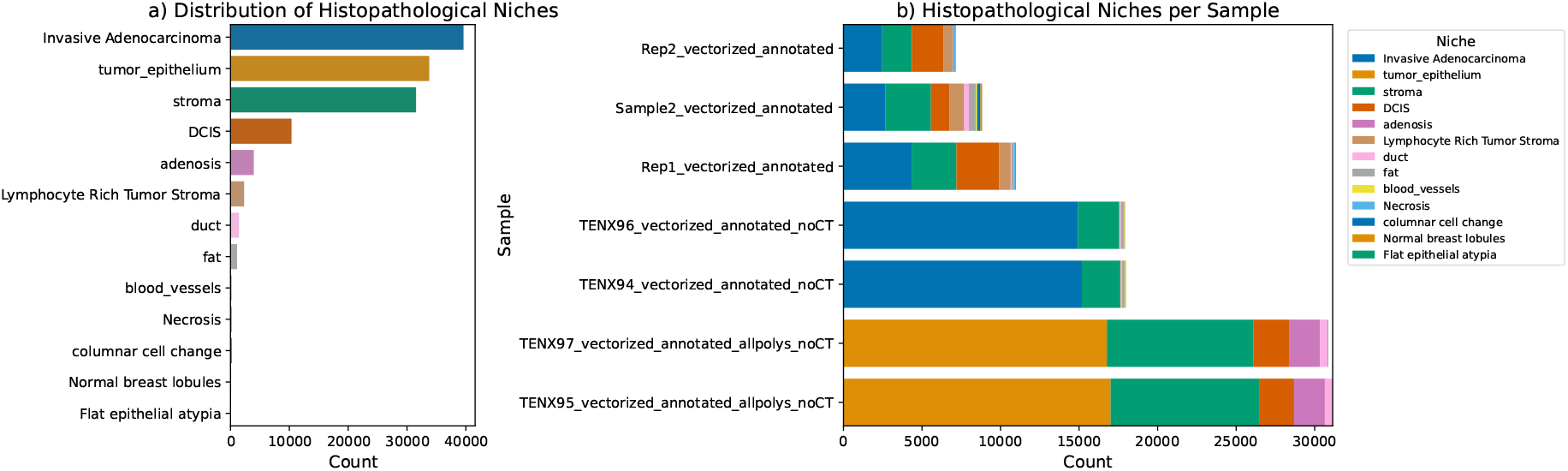
Data exploration for the breast dataset. **(a)** Overall niche distribution excluding *NOANNOT* patches. The breast cancer samples show tumor microenvironment complexity with distinct tumor regions, stromal compartments, immune infiltration zones, and adipose tissue. **(b)** Niche distribution per sample reveals tumor heterogeneity (7-10 distinct niches per sample), with variable immune and stromal content reflecting different cancer subtypes and stages. No cell-type composition labels are available for certain samples (marked with “*” in Supplementary Table 1).

#### Niche heterogeneity and task complexity

The thymus dataset exhibits the highest per-sample niche diversity (10-11 niches/sample), reflecting its highly structured developmental organization. The breast dataset shows moderate heterogeneity (7-10 niches/sample) driven by tumor microenvironment complexity. The lung dataset displays the widest range (2-13 niches/sample), with heterogeneity correlating with disease severity. This variation in niche complexity directly impacts the difficulty of cross-modal fusion: higher heterogeneity increases the number of distinct morphology-expression relationships the model must learn, making synergy-aware integration progressively more valuable.

### 3 Training data for pretrained encoders

#### Nicheformer (gene expression encoder)

We use the pretrained Nicheformer encoder described in Tejada-Lapuerta et al. [8]. Nicheformer is pretrained on SpatialCorpus-110M, a large-scale collection comprising approximately 57.06 million dissociated single-cell transcriptomes and 53.83 million imagebased spatially resolved cells across 73 human and mouse tissues. The SpatialCorpus-110M dataset is publicly available via Hugging Face at https://huggingface.co/datasets/theislab/SpatialCorpus-110M. We use the authors’ released pretrained model checkpoint, provided through Mendeley Data at https://data.mendeley.com/preview/87gm9hrgm8?a=d95a6dde-e054-4245-a7eb-0522d6ea7dff. Unless explicitly stated otherwise (e.g., for fine-tuning experiments), Nicheformer is treated as a frozen encoder.

#### UNI-2 (image encoder)

We use UNI-2 (UNI2-h) as described in Chen et al. [5]. UNI-2 is pretrained using self-supervised learning on a large corpus of histopathology image tiles (>200M tiles) derived from over 300k-350k whole-slide images sourced from internal clinical archives at Mass General Brigham. As detailed in the corresponding publication, the underlying pretraining data are not fully publicly released due to institutional and regulatory constraints, and access to curated datasets is evaluated on a case-by-case basis. We use the publicly released `UNI2-h` model checkpoint available at https://huggingface.co/MahmoodLab/UNI2-h. Unless explicitly noted, UNI-2 is used as a frozen image encoder.

### 4 Fusion method details

In the main text, we compare the performance of different multimodal fusion methods. We give short descriptions of how we implement these approaches below. As detailed in the Methods, most approaches learn modality-specific fusion mappings *g*_1_ and *g*_2_ and produce projected representations 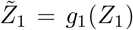 and 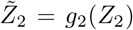. Unless stated otherwise (e.g., for specific baselines like Concatenation or Random), downstream evaluation concatenates 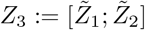 and trains a linear probe.

#### Barlow Twins [44]

This approach uses redundancy reduction by computing the cross-correlation matrix between modality-specific features and regularizing it to be close to the identity matrix. This enforces invariance across modalities (diagonal terms) while discouraging trivial feature redundancy within the projected space (off-diagonal terms). Hyperparameters: *λ* = 5 × 10^−3^ (off-diagonal weight), projection dimension = 256. Architecture: per modality `Linear(input dim, 256, bias=False)` followed by batch normalization without affine parameters. We use a linear projector variant in this frozen encoder benchmark to isolate objective-level differences; all spectral analysis statements in the main text apply to this linear-projector setting. This linear architecture ensures consistency with the spectral theory, as Barlow Twins reduces to trace maximization in the linear regime [27].

#### BYOL [46]

Bootstrap Your Own Latent (BYOL) uses asymmetric regression with a stop-gradient target. In our bimodal setting, we maintain an *online* branch and a *target* branch per modality: each modality has an online projector and predictor, and a target projector updated by exponential moving average (EMA). Given paired samples, the online prediction from modality #1 is regressed onto the (stopgradient) target of modality #2, and vice versa. We use EMA momentum *τ*_ema_ = 0.996. Architecture: per modality, projector `Linear(inpu_dim, 256`); predictor `Linear(256, 128) → LayerNorm →GELU → Linear(128, 256)`.

#### CCA

Linear Canonical Correlation Analysis (CCA) finds linear projections that maximize cross-modal correlation under whitening constraints. On frozen representations, the solution is given by an singular value decomposition (SVD)/eigendecomposition of the *whitened* cross-covariance (equivalently, a generalized eigenvalue problem). We use ridge regularization *r*_1_ = *r*_2_ = 10^−4^ for numerical stability and set projection dimension = 256.

#### CoMM [34]

Contrastive Multimodal (CoMM) implements partial information decomposition loss with explicit synergy terms. It uses transformer-based fusion optimizing for complementary information rather than mere alignment. Hyperparameters: temperature *T* = 0.07. Architecture: transformer-based fusion with cross-attention (1 layer, 8 heads, embedding dimension = 256), with linear adapters projecting each modality to the shared dimension, followed by a 2-layer ReLU projection head.

#### Concatenation Baseline

Simple baseline that concatenates the frozen unimodal embeddings from the gene expression encoder and the image encoder without any learned fusion. There are no trainable parameters.

#### DCCA [65]

Deep Canonical Correlation Analysis (DCCA) replaces the linear CCA projections with neural networks and optimizes the CCA objective on their outputs. We use batch normalization and sigmoid activations in our implementation. Hyperparameters: learning rate = 10^−4^, regularization *r*_1_ = *r*_2_ = 10^−3^. Architecture: 3-layer MLP per modality `Linear(input_dim, 1024) → Sigmoid → BatchNorm1d → Linear(1024, 1024) → Sigmoid → BatchNorm1d → Linear(1024, 256)`.

#### DIM (Neural) [45]

Deep InfoMax (DIM) uses a parameterized critic to distinguish matched crossmodal pairs from mismatched pairs, yielding a neural lower bound proxy to mutual information. We implement a critic that scores concatenated embedding pairs and train it with matched pairs as positives and within-batch mismatches as negatives. Hyperparameters: temperature *T* = 0.1, normalized embeddings. Architecture: 3-layer encoder per modality `Linear(input_dim, hidden_dim) → LayerNorm → GELU → Linear(hidden dim, 256) → LayerNorm → GELU → Linear(256, 256`), with an MLP discriminator `Linear(512, 256) → GELU → Linear(256, 1)` that scores concatenated embedding pairs.

#### Random Baseline

Negative control using random Gaussian vectors that are the same dimensionality as the multimodal embeddings, sampled from a standard normal distribution 𝒩 (0, *I*).

#### SimCLR [36]

Simple Contrastive Learning of Representations (SimCLR) is a contrastive learning framework that maximizes agreement between different views using normalized temperature-scaled cross entropy loss (NT-Xent), where positive pairs are diagonal elements of the similarity matrix and negative pairs are off-diagonal. Hyperparameters: temperature *τ* = 0.1, batch size = 256 (override of the default batch size). Architecture: 2-layer MLP per modality `Linear(input_dim, 256) → LayerNorm → GELU → Linear(256, 256)`.

#### SimSiam [47]

Simple Siamese (SimSiam) networks use a predictor network to prevent collapse and demonstrate that stop-gradient is sufficient for learning meaningful representations without momentum encoders or negative samples. This approach uses symmetric cosine similarity loss with stop-gradient on one branch. Architecture: 3-layer encoder per modality `Linear(input_dim, 512) → LayerNorm → GELU → Linear(512, 256) → LayerNorm → GELU → Linear(256, 256)`, followed by a 2-layer predictor `Linear(256, 128) → BatchNorm1d → ReLU → Linear(128, 256)`.

#### VICReg [35]

Variance-Invariance-Covariance Regularization (VICReg) combines (i) an invariance term that matches paired projections, (ii) a variance term that prevents collapse, and (iii) a covariance term that decorrelates feature coordinates within each modality’s projected space. Hyperparameters: similarity coefficient = 25, variance coefficient = 25, covariance coefficient = 1, stability epsilon = 10^−4^. Architecture: per modality `Linear(input_dim, 256`) followed by batch normalization. This linear architecture ensures consistency with the spectral theory, as VICReg reduces to trace maximization in the linear regime [27].

### 5 General method implementation details

#### Training protocol

Unless otherwise noted, all fusion methods are trained with batch size 32 using AdamW (learning rate 10^−4^, weight decay 10^−5^), up to 100 epochs with early stopping (patience 10) on validation loss, mixed precision, and gradient clipping (norm 1.0). Methods that require larger batches for stable contrastive estimation use method-specific batch sizes (e.g., SimCLR uses batch size 256). Unsupervised methods (CoMM, SimCLR, Barlow Twins, VICReg, BYOL, SimSiam, and DIM) use the full dataset; CCA and DCCA use closed-form or iterative optimization. All experiments use mixed precision training and gradient clipping at norm set to 1.0.

#### Evaluation protocol

Linear probes are implemented using `scikit-learn: LogisticRegression` for classification tasks and `RidgeCV` for regression tasks. `RidgeCV` uses cross-validation to select the optimal regularization parameter from `alphas=np.logspace(-3, 3, 10)` with *R*^2^ scoring. All probes use 5-fold cross-validation or train-test splits depending on the evaluation strategy.

### 6 Extended spatial analysis

In the main text, we showed that fusion gains (as measured by SIS) vary with spatial complexity in the lung dataset. Here, we replicate the same spatial neighborhood evaluation on the breast (Xenium; high-resolution) and thymus (Visium; spot-based) datasets to test whether these trends generalize across resolution regimes. Our theoretical framing predicts that when cross-modal correspondence is tight at high resolution, multimodal gains are primarily redundancy-limited, whereas resolution mismatch or spatial ambiguity can increase the value of integration-oriented interfaces.

#### Breast (high-resolution Xenium)

The breast dataset is characterized by high cellular density and patch-level correspondence between morphology and gene expression. Neighbor niche prediction is consistent with a redundancy-dominated regime: concatenation performs comparably to, or slightly better than, learned fusion interfaces across distance bins (Supplementary Fig. 4a). SIS is near zero or negative at short range (*<* 10 *µ*m; Supplementary Fig. 4b), and increases with distance as local correspondence weakens and predictions become more ambiguous.

#### Thymus (spot-based Visium; resolution mismatch)

The thymus dataset uses Visium (55 *µ*m spots), inducing a scale mismatch between cell-level morphology and spot-level expression aggregates. In this regime, CoMM provides a more consistent advantage over concatenation at intermediate distances (Supplementary Fig. 4e), and SIS remains positive over a broader range of distances (Supplementary Fig. 4f). These results are consistent with the hypothesis that integration-oriented interfaces can be more beneficial when correspondence is imperfect due to measurement resolution.

#### Note on consistency metrics

Across both datasets, spatial and niche consistency (Supplementary Fig. 4c,d,g,h) remain comparable to strong unimodal baselines, indicating that fusion interfaces do not trivially destroy neighborhood geometry. For the lung dataset, complete consistency results are reported in Supplementary Note 7. In this diagnostic, low or negative SIS should be interpreted as limited incremental information gain under the chosen probe class, rather than a loss of intrinsic embedding structure.

**Supplementary Figure 4.**
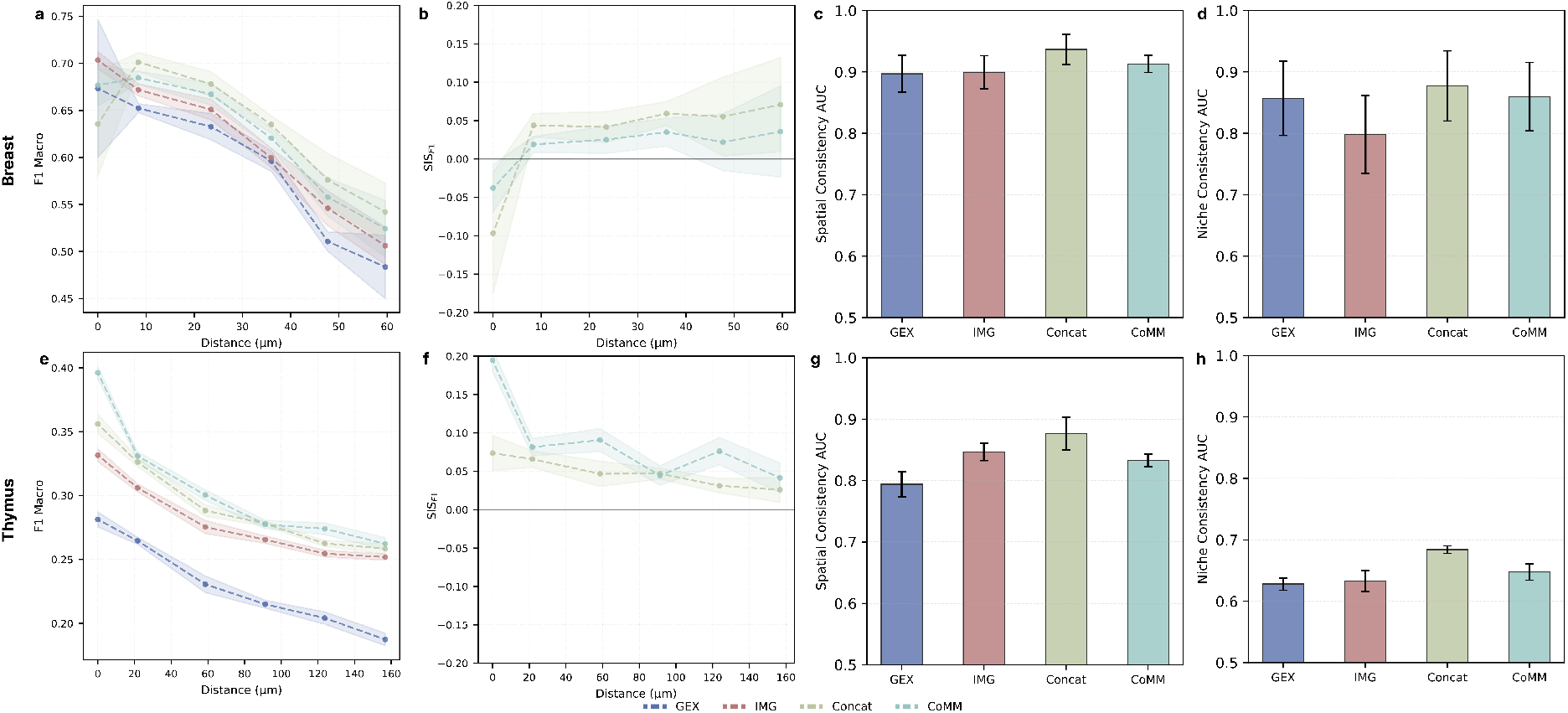
Breast (redundancy-dominated; panels a-d): **(a)** Macro F1 for neighbor niche prediction across distance bins; **(b)** SIS computed from F1; **(c)** Spatial consistency (area under the curve; AUC); **(d)** Niche consistency (AUC). **Thymus (resolution-mismatched; panels e-h): (e)** Macro F1 for neighbor niche prediction; **(f)** SIS computed from F1; **(g)** Spatial consistency (AUC); **(h)** Niche consistency (AUC). Shaded regions and error bars denote variability across folds.

### 7 Complete results on spatial and niche consistency

The main text focuses on *fusion gain* via SIS. Here, we report complementary consistency diagnostics that quantify whether a fusion interface preserves the intrinsic geometry already present in strong unimodal encoders. Concretely, we test whether a lightweight linear decoder applied to pairwise embedding differences can reliably recover (i) *spatial locality* and (ii) *semantic niche identity* from frozen embeddings. These diagnostics serve two purposes: (1) they verify that interfaces do not destroy the neighborhood structure needed for retrieval-style use cases, and (2) they help interpret low SIS values as *limited incremental contribution* rather than a collapse of representation structure.

#### Diagnostic #1: spatial consistency (locality preservation)

We evaluate whether patches that are spatially close on the same slide are embedded closer than spatially distant patches. Positive pairs are 5-nearest neighbors within a slide (approximately *<* 10 *µ*m separation); negative pairs are pairs beyond the 80th percentile within-slide distance (approximately > 50 *µ*m), yielding *N* = 120,832 pairs. We train a logistic regression probe on absolute embedding differences to predict the pair label and report classification and retrieval metrics. Results are shown in Supplementary Table 2.

**Supplementary Table 2.** Spatial consistency metrics for the lung dataset. Reported values are mean ± standard deviation over 5-fold GroupKFold cross-validation on *N* = 120,832 patch pairs. Positive pairs are 5-nearest neighbors within the same slide; negative pairs are pairs with distance above the 80th percentile. Metrics include macro F1, accuracy (ACC), area under the curve (AUC), and average precision (AP). IMG = imaging, GEX = gene expression, and Concat = Concatenation.

| Method | F1 | ACC | AUC | AP |
| --- | --- | --- | --- | --- |
| Unimodal IMG | 0.844±0.014 | 0.785±0.012 | 0.853±0.029 | 0.936±0.007 |
| Unimodal GEX | 0.860±0.016 | 0.804±0.020 | 0.867±0.031 | 0.943±0.009 |
| Multimodal Concat | 0.881±0.011 | 0.834±0.015 | 0.903±0.024 | 0.960±0.006 |
| Multimodal CoMM | 0.857±0.020 | 0.801±0.025 | 0.870±0.032 | 0.945±0.009 |

**Supplementary Table 3.** Niche consistency metrics for the lung dataset. Reported values are mean± standard deviation over 5-fold GroupKFold cross-validation on *N* = 200,000 patch pairs. Positive pairs share the same niche label; negative pairs correspond to different niche labels. Metrics include macro F1, accuracy (ACC), area under the curve (AUC), and average precision (AP). IMG = imaging, GEX = gene expression, and Concat = Concatenation.

| Method | F1 | ACC | AUC | AP |
| --- | --- | --- | --- | --- |
| Unimodal IMG | 0.758±0.060 | 0.757±0.055 | 0.833±0.057 | 0.811±0.066 |
| Unimodal GEX | 0.760±0.043 | 0.756±0.039 | 0.830±0.039 | 0.802±0.041 |
| Multimodal Concat | 0.796±0.046 | 0.794±0.044 | 0.872±0.040 | 0.851±0.046 |
| Multimodal CoMM | 0.776±0.047 | 0.774±0.044 | 0.850±0.044 | 0.829±0.050 |

#### Diagnostic #2: niche consistency (semantic neighborhood preservation)

We evaluate whether patches sharing the same niche label are embedded closer than patches from different niches. We sample *N* = 200,000 pairs balanced between same-niche (positive) and different-niche (negative) examples. As

#### Interpreting the distance baseline

The distance baseline evaluates retrieval directly from the frozen embedding geometry: we first compute Euclidean distances between patch embeddings using − *d*(*e*_*i*_, *e*_*j*_) as a score (smaller distance ⇒ higher score), then we compute the area under the curve (AUC) or average precision (AP) without learning a task-specific probe. The learned probe improves over this baseline because it can reweight embedding dimensions to match the pairwise label, while remaining a lightweight linear decoder. This separation clarifies what changes: the diagnostic is not about updating embeddings, but about how well locality/semantics are *decodable* from the frozen geometry with minimal supervision.

#### Key findings

Across both diagnostics, multimodal interfaces preserve spatial and semantic neighborhood structure at least as well as the strongest unimodal baselines (Supplementary Tables 2-3). In particular, concatenation attains the highest AUC in both spatial consistency (AUC = 0.903) and niche consistency (AUC = 0.872), consistent with the redundancy-dominated regime in the lung dataset: when each modality already carries a coherent neighborhood structure, simple aggregation preserves both without enforcing cross-modal redundancy reduction. These results complement SIS: they show that low SIS does not imply unusable embeddings; rather, many interfaces preserve geometry well but do not yield additional *incremental* probe-accessible information beyond the strongest unimodal baseline.

**Supplementary Figure 5.**
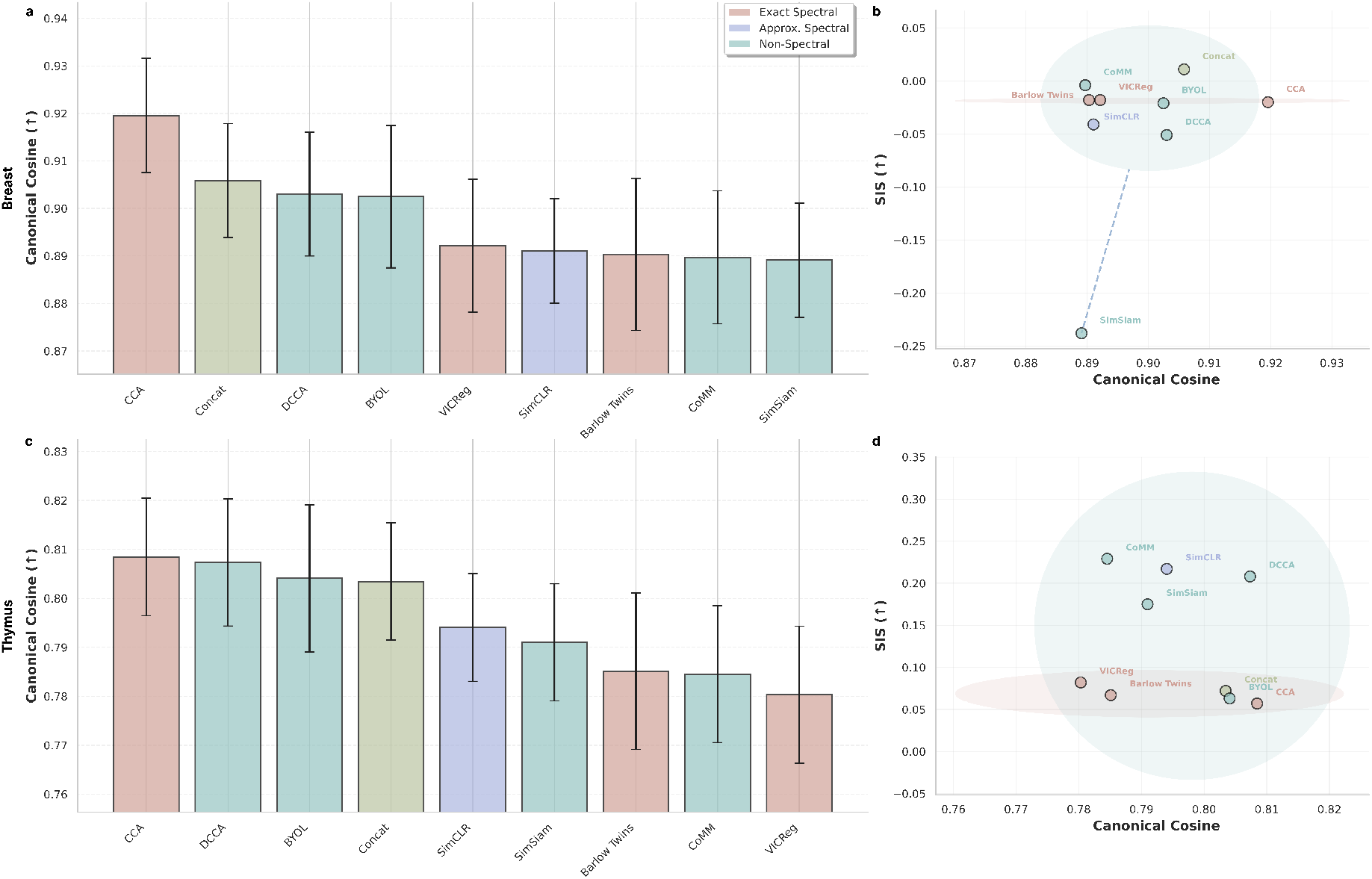
Spectral ceiling and synergy across datasets. Breast (top): **(a)** canonical cosine quantifying convergence to the linear subspace recovered by spectral solutions; **(b)** alignment-synergy landscape (SIS versus canonical cosine). **Thymus (bottom): (c)** canonical cosine; **(d)** SIS versus canonical cosine. Error bars and shaded regions denote variability across folds.

### 8 Spectral ceiling and alignment-synergy across datasets datasets

We repeat the same spectral diagnostics used in the lung dataset to assess whether the alignment-synergy trade-off generalizes to breast and thymus (Supplementary Fig. 5). In the breast dataset (Supplementary Fig. 5a,b), methods concentrate at high canonical cosine with SIS near zero, consistent with a regime where recoverable cross-modal signal is largely captured by linear redundancy. In the thymus dataset (Supplementary Fig. 5c,d), canonical cosine values are lower overall, reflecting more challenging correspondence under spot-level expression and higher-resolution imaging. Notably, several integration-oriented methods attain higher SIS while exhibiting canonical cosine comparable to spectral baselines, consistent with accessing task-relevant structure not explained by the linear spectral solution.

#### Description of metrics

To quantify convergence to the theoretical spectral solution, we compute canonical cosine following standard numerical linear algebra protocols [49, 66]. We compute principal angles between the column space of the learned projections and the top-*K* singular-vector subspace from the cross-covariance SVD, then report the mean cosine of these angles. Canonical cosine ranges from 0 to 1, with 1 indicating perfect subspace alignment. Unless stated otherwise, canonical cosine is computed on *N* = 1000 held-out test patches with effective dimensionality *K* = 5 determined by parallel analysis using *L* = 100 permutations.

### 9 Scaling experiment implementation details

#### Data splitting strategy

For the scaling analysis in the main text, we first create a fixed 80/20 train/test split from the full dataset. The test set remains fixed across all scaling experiments to ensure fair comparison and prevent data leakage. We then create partitions of the training set at log-scale fractions: 1%, 3.16%, 10%, 31.6%, and 100% of the training data. Each partition is sampled randomly from the training set without replacement.

#### Fine-tuning protocol

For each partition, we fine-tune the pretrained Nicheformer gene expression encoder on the subset of training data. Fine-tuning uses an AdamW optimizer with a cosine annealing learning rate schedule and warmup. Hyperparameters vary by dataset and are determined via a sweep: for the lung dataset, we use learning rate = 2 × 10^−5^, weight decay = 5 × 10^−4^, batch size = 16 with gradient accumulation, and up to 100 epochs with early stopping (patience 50 epochs) based on validation loss. The validation set is created as a 10% split from each training partition. All fine-tuned models are evaluated on the same held-out test set using 5-fold cross-validation to compute performance metrics and SIS scores. All performance metrics (macro F1) are evaluated on held-out test sets. SIS values are derived post hoc from trained models.

#### Multimodal evaluation

After fine-tuning the gene expression encoder, we evaluate multimodal fusion methods (CoMM and Concatenation) using the fine-tuned encoder and frozen image encoder. This allows us to assess whether multimodal integration provides additional value beyond the adapted unimodal baseline. The multimodal models are trained using the same fusion protocol as described in Supplementary Note 5, with embeddings extracted from the fine-tuned gene expression encoder.

### 10 Scaling: extended fine-tuning analysis across datasets

We extend the main text scaling experiment by repeating the unimodal fine-tuning analysis on the breast and thymus datasets (Supplementary Fig. 6). Here, the goal is to test whether sample-efficient unimodal adaptation persists across resolution regimes, and assess whether multimodal fusion provides additional gains once a strong unimodal encoder is adapted.

**Supplementary Figure 6.**
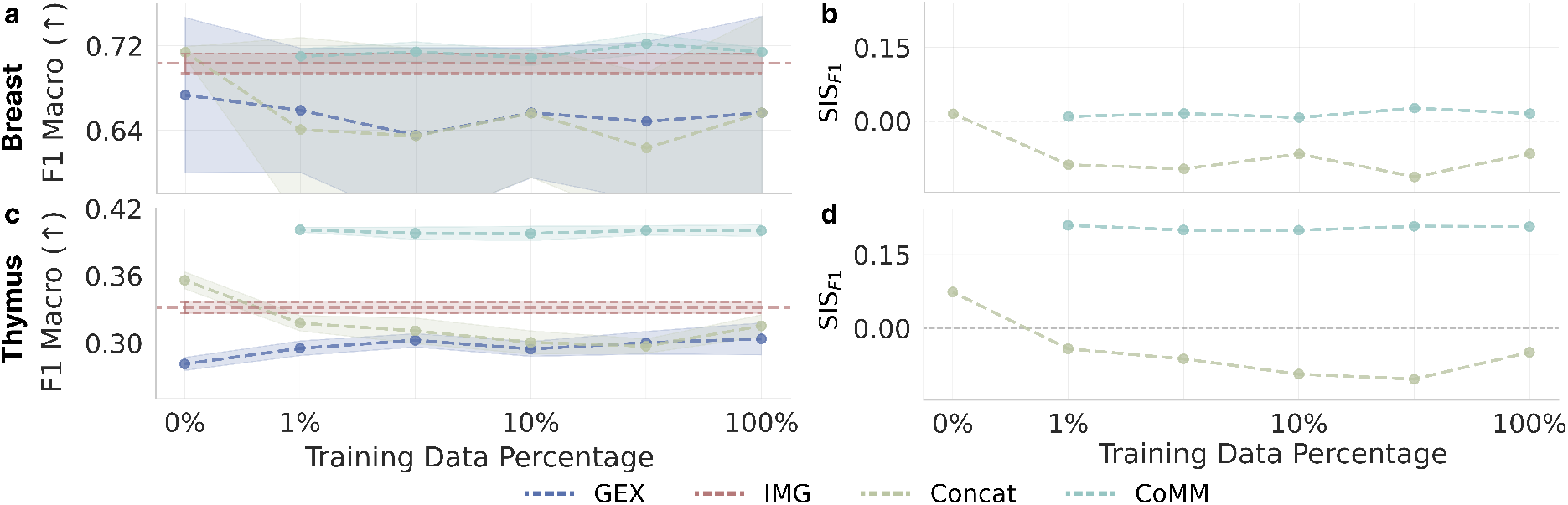
Extended scaling analysis. Breast (up): **(a)** Macro F1 for niche classification versus fraction of data used for fine-tuning; and **(b)** corresponding SIS as a function of the fraction of data used for fine-tuning. **Thymus (down): (c)** Macro F1 versus fraction of fine-tuning fraction; and **(d)** corresponding SIS.

#### Thymus (slower unimodal catch-up under resolution mismatch)

On the thymus dataset, unimodal gene expression (GEX) fine-tuning shows modest gains, while CoMM remains a strong but constant baseline. SIS remains positive for CoMM, while it degrades for Concatenation.

#### Breast (rapid unimodal saturation)

On the breast dataset, changes from unimodal GEX finetuning are within experimental variation. Its performance remains high overall, while CoMM maintains a slightly positive SIS and we observe degradation for the Concatenation baseline.

#### Takeaway

Across datasets, these scaling results support a practical design principle for compositional foundation models: when a single modality can be efficiently adapted to the task, multimodal fusion often yields limited incremental gain under a linear probe. Conversely, resolution mismatch and spatial ambiguity are the regimes where integration-oriented interfaces are most likely to help, particularly at limited paired-data budgets.

